# Nanopore sequencing for fast determination of plasmids, phages, virulence markers, and antimicrobial resistance genes in Shiga toxin-producing *Escherichia coli*

**DOI:** 10.1101/571364

**Authors:** Narjol Gonzalez-Escalona, Marc A. Allard, Eric W. Brown, Shashi Sharma, Maria Hoffmann

## Abstract

Whole genome sequencing can provide essential public health information. However, it is now known that widely used short-read methods have the potential to miss some randomly-distributed segments of genomes. This can prevent phages, plasmids, and virulence factors from being detected or properly identified. Here, we compared assemblies of three complete STEC O26:H11 genomes from two different sequence types (ST21 and 29), each acquired using the MiSeq-Nextera XT, MinION nanopore-based sequencing, and Pacific Biosciences (PacBio) sequencing. Each closed genome consisted of a single chromosome, approximately 5.7 Mb for CFSAN027343, 5.6 Mb for CFSAN027346, and 5.4 MB for CFSAN027350. However, short-read WGS using MiSeq-Nextera failed to identify some virulence genes in plasmids and on the chromosome, both of which were detected using the long-read platforms. Results from long-read MinION and PacBio allowed us to identify differences in plasmid content: a single 88 kb plasmid in CFSAN027343; a 157kb plasmid in CFSAN027350; and two plasmids in CFSAN027346 (one 95 Kb, one 72 Kb). These data enabled rapid characterization of the virulome, detection of antimicrobial genes, and composition/location of Stx phages. Taken together, positive correlations between the two long-read methods for determining plasmids, virulome, antimicrobial resistance genes, and phage composition support MinION sequencing as one accurate and economical option for closing STEC genomes and identifying specific virulence markers.

## Introduction

Whole genome sequencing is an essential tool for characterizing and tracking pathogenic bacteria that may have contaminated the food supply as well as for identifying whether those bacteria carry virulence factors that could pose serious threats to public health (1,2). Closed bacterial genomes provide: 1) high-quality reference material that supports pathogen source tracking during a foodborne outbreak investigation, 2) important clues about the long-term evolution of enteric pathogens, 3) key insights into mechanisms and transmission of mobile elements conferring antimicrobial resistance, and 4) critical information about the contribution of DNA modifications on pathogenesis (2–9). These details are particularly important for understanding hemorrhagic pathogens such as *E. coli* O26:H11/- strains, which cause significant human morbidity and mortality worldwide (31,33,35,43). The genomes of O26:H11/- are very complex, containing many virulence genes, insertion sequences, phages, and plasmids; consequently, strains of the same lineage can possess significantly different content (7,9,35),(33,36–38). Missing the presence of some of these elements during an investigation can have large impacts on human health (e.g. *hlyA* gene in EHECs). Thus, it really matters that we have reproducible WGS systems for fully capturing and sequencing these elements.

Long-read sequencing platforms afford one solution to this challenge. Some systems such as Pacific Biosciences (PacBio) Sequencers *RSII* or Sequel (https://www.pacb.com/products-and-services/pacbio-systems/), use single-molecule real-time (SMRT) sequencing technology that allow for real-time observation of DNA synthesis through zero-mode waveguides (ZMWs) and phospho-linked nucleotides (11,12). While comprehensive in their ability to capture entire genomes, extraneous elements included, these systems often require significant investments in machinery, space, and laboratory expertise, all of which may be obstacles to routine use. These systems also require significant quantities of DNA *(i.e.,* 5 μg), require a more substantial preparatory time (i.e., 8 hr DNA sequencing library protocols, and produce average read lengths of about 11Kb, although reads of greater than 50 kb were possible in our laboratory at the time of this study.

Alternative sequencing platforms based on nanopore technology may be able to provide high-quality libraries from long reads and produce closed bacterial genomes, while also offering several other advantages in portability and affordability. These systems are much less expensive, take little laboratory space, and can even be taken into the field for on-site sequencing. The MinION nanopore (Oxford Nanopore, Oxford, UK) system, as one example, comprises a palm-sized unit able to detect changes in ionic current when DNA or RNA passes through the nanopores, whereupon those changes are translated into base calls. Researchers have already used nanopore sequencing on plants, yeasts, viruses, and to perform *de novo* bacterial assembly (14,16,21). Other applications have included rapid identification of viral pathogens (18,22), metagenomics (22–24), detection of antimicrobial resistance genes (25,26), comparative RNA expression levels in diverse cells (27), and detecting DNA methylation patterns (28). Since the initial MinION Access Programme (MAP), this technology has been refined several times (15). At least three nanopore-based sequencing tools are currently in production as of autumn 2018 (13–16).

As MinION does not require a size selection step prior to sequencing, it can obtain longer reads than other platforms (13–15, 17–19). Moreover, the system also allows for “quasi” real-time sequencing approaches (20). The accuracy of nanopore sequencing is reported to be 90% in general, with some researchers reporting 99.96% accuracy only after read polishing (http://simpsonlab.github.io/2016/08/23/R9).

Two of the main subgroups within O26:H11/- are the enteropathogenic (EPEC), which carry the locus of enterocyte effacement (LEE = locus of enterocyte effacement) causing mild diarrhea, and enterohemorrhagic (EHEC), which carry either stx1 and/or sxt2 genes in addition to the LEE, and are associated with more severe illnesses, such as hemorrhagic colitis (HC) and hemolytic uremia syndrome (HUS) (34,44). A new clone of O26:H11/H- belonging to sequence type 29 (ST29) and that had a specific virulome (*stx2*_a_+, *eae*+, plasmid gene profile *ehxA*+, *etpD*+) has been recently found distributed all over Europe (35). Serotype O26:H11/H- can be divided, based on MLST, into several STs, with ST21 and ST29 associated with disease in humans (7). We have selected three STEC O26:H11 strains with very complex genomes belonging to different STs (21 and 29), and isolated in different years, places and sources (clinical and environmental). This makes them ideal strains to compare the three WGS capabilities.

In this study, we compared the sequencing capabilities of the MinION with that of the PacBio RS II and MiSeq by sequencing 3 STEC O26:H11 strains (see below) (35) using MinION and a genome library prepared using two different library kits *(i.e.,* 1D ligation kit and rapid sequencing kit). We then compared the results obtained by this sequencing platform (MinION). We also tested two different assembly pipelines for the *de novo* assembly of these genomes. Finally, we compared the results of each technology to assess their capacity for detecting virulence and antimicrobial genes, Shiga toxin phages, plasmid presence, genome synteny, and phylogenetic analysis.

## RESULTS

The 3 STEC strains employed here are listed in Table 1, along with their sources and confirmed STs. These genomes were sequenced on each of the sequencing systems, and three main differences were documented among those systems: number of contigs, ease of assembly, and detection of important genomic markers.

**Table 1.** Summary of the characteristics of the 3 STEC O26:H11 strains sequenced in this study.

| strain | CFSAN No. | Serotype<sup>a</sup> | ST<sup>b</sup> | Date | location | source |
| --- | --- | --- | --- | --- | --- | --- |
| 99.085 | CFSAN027343 | O26:H11 | 21 | 1999 | Argentina | clinical |
| 99.1773 | CFSAN027346 | O26:H11 | 21 | 1999 | USA | clinical |
| 12.1843 | CFSAN027350 | O26:H11 | 29 | 2012 | USA | environmental |

<sup>a</sup>These results were confirmed with *in silico* serotyping.
<sup>b</sup>Determined by *in silico* MLST.

### MiSeq sequencing and assembly

When these STECs were sequenced using MiSeq (Illumina) and assembled *de novo* using Nextera XT, their genomes assembled into 234 to 321 contigs (Table 2), which is within the expected range for STECs; results from other closed STECs typically >250 contigs (based on our own and other researcher’s observations; data available from NCBI). *in silico* MLST identified the samples correctly as belonging to ST21 (CFSAN027343 and CFSAN027346), and ST29 (CFSAN027350). *in silico* serotyping confirmed that these strains were all serotype O26:H11.

**Table 2.** MiSeq assembly statistics for the 3 O26:H11 STECs.

| CFSAN No. | Contigs<br>MiSeq | No.<br>of reads | Q>30<br>Reads<br>size | Total<br>bases<br>(bp) | N50 | Total<br>genome<br>(bp) | Average<br>coverage<br>X |
| --- | --- | --- | --- | --- | --- | --- | --- |
| CFSAN027346 | 274 | 2.14E+06 | 194 | 4.17E+08 | 100,594 | 5.42E+06 | 77 |
| CFSAN027343 | 321 | 2.56E+06 | 188 | 4.81E+08 | 93,611 | 5.41E+06 | 89 |
| CFSAN027350 | 234 | 2.45E+06 | 221 | 5.42E+08 | 99,163 | 5.28E+06 | 103 |

*de novo* assembly of the 3 genomes resulted in assemblies that ranged from 5.2 (CFSAN027350) to 5.4 Mb (the other two genomes) (Table 2). As stated earlier, it is notable that *de novo* assemblies of short-read sequencing technology produced an elevated number of contigs resulting in fractioned assemblies.

### MinION sequencing and DNA sequencing library approaches

We sequenced the same three STECs using the nanopore-based MinION device. Two strains were sequenced in triplicate (CFSAN027350 and CFSAN027346); CFSAN027343 was sequenced twice. We used two different DNA library approaches: the 1D ligation kit (SQK-LSK108, expected to produce more output and longer reads) and the rapid sequencing kit (SQK-RAD002, expected to produce lower output and smaller reads, but with a much simpler and straightforward procedure for preparing the DNA library).

Table 3 confirms that the 1D ligation kit produced larger outputs than the rapid sequencing kit. Even though the DNA extraction method was the same for each strain, the sequencing output was different, varying from 1 Gb (CFSAN027350a) to 8 Gb (CFSAN027343a). DNA libraries output also varied between replicates (with a higher percentage of reads > 5 kb in the case of CFSAN027346b with 355,653 reads above 5 kb long compared to the same replicate library CFSAN027346a with 139,883 reads above 5 kb (results not shown). As expected, sequencing runs prepared with the rapid kit produced lower output, with an average of 0.71 Gb (Table 3).

**Table 3.**
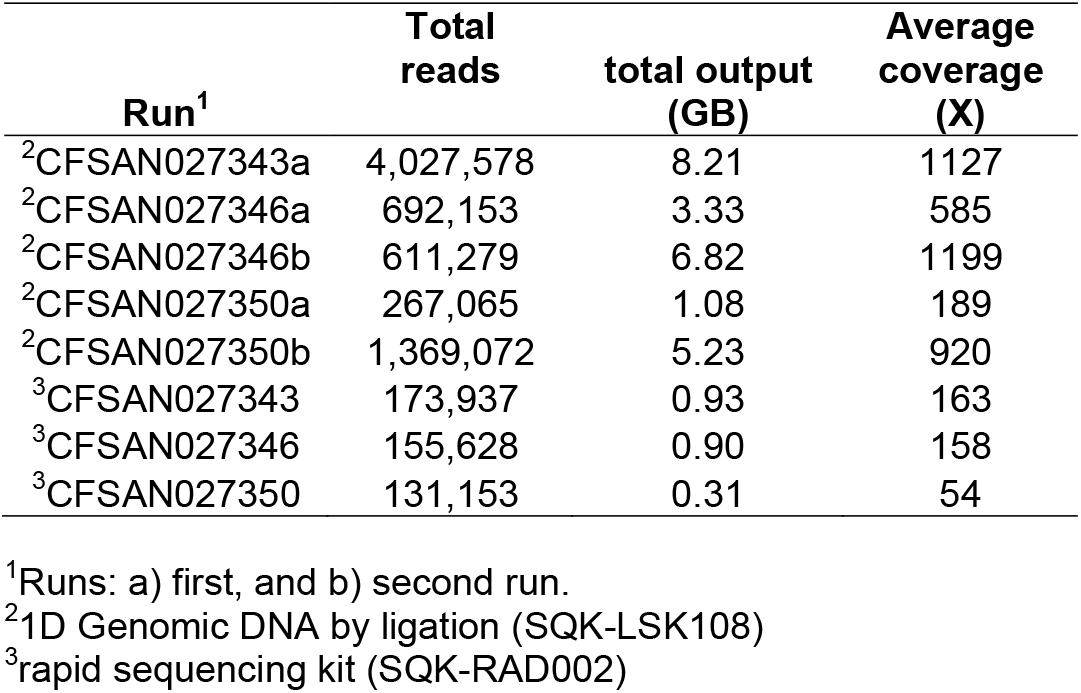
STECs MinION sequencing output statistics by replicate and library kit.

Nevertheless, after running our CANU assembly, we were able to produce a closed bacterial genome for each strain, including the chromosome and plasmid(s) (Table 4). Although there were still variations in chromosome sizes among the replicates, we found overall agreement that CFSAN027343 and CFSAN027350 each contained a chromosome of ∼ 5,6 Mb with a single plasmid while CFSAN027346 contained two plasmids. Strikingly, the plasmid identified in CFSAN027350 was 155 kb, larger than any previously described for O26:H11 STECs. We will describe this plasmid in more detail later in this section.

**Table 4.** Assembly statistics MinION.

| Sample<sup>1</sup> | Chromosome(s)<br>contig(s) | plasmids | Chromosome<br>size (bp) | plasmid(s)<br>size(s) (bp) |
| --- | --- | --- | --- | --- |
| <sup>2</sup>CFSAN027343a | 1 | 1 | 5,688,712 | 88,561 |
| <sup>3</sup>CFSAN027343 | 1 | 1 | 5,688,145 | 88,702 |
| <sup>2</sup>CFSAN027346a | 1 | 2 | 5,588,947 | 1 (95,599);<br>2 (72,940) |
| <sup>2</sup>CFSAN027346b | 1 | 2 | 5,592,589 | 1 (95,821);<br>2 (72,972) |
| <sup>3</sup>CFSAN027346 | 1 | 2 | 5,592,692 | 1 (95,696);<br>2 (72,950) |
| <sup>2</sup>CFSAN027350a | 1 | 1 | 5,422,984 | 157,276 |
| <sup>2</sup>CFSAN027350b | 3 | 1 | 5,448,646 | 157,340 |
| <sup>3</sup>CFSAN027350 | 15 | 1 | 5,451,905 | 157,300 |

<sup>1</sup>Runs: a) first, and b) second run.
<sup>2</sup>1D Genomic DNA by ligation (SQK-LSK108)
<sup>3</sup>rapid sequencing kit (SQK-RAD002)

### PacBio sequencing

We next sequenced our three STECs using the PacBio *RSII* system with a 20-kb insert library protocol. After the library protocol was completed, we sequenced it on three SMRT cells (Table 5). Although the same preparation was used for each isolate, slight differences were noted in the output of raw data among different SMRT cells, (Table 5). The total output for CFSAN027343, CFSAN027346, and CFSAN027350 was 2.3, 2.84. and 2.9 Gb, respectively. The small discrepancy is common since the SMRT cells were loaded with a different binding complex for each strain. The average read of insert lengths for each individual strain CFSAN027343, CFSAN027346, and CFSAN027350 was 10,011 bp, 10,804 bp, and 11,632 bp, respectively.

**Table 5.** PacB¡o sequencing output for each SMRT cell

| Strains | SMRT<br>Cell# | Total<br>Reads | Total<br>Output<br>(GB) | Average Read<br>of Insert length |
| --- | --- | --- | --- | --- |
| CFSAN027343 | 1 | 79,102 | 0.79 | 9,965 |
|  | 2 | 77,649 | 0.80 | 10,326 |
|  | 3 | 75,856 | 0.74 | 9,743 |
| CFSAN027346 | 1 | 91,744 | 1.03 | 11,252 |
|  | 2 | 82,672 | 0.88 | 10,677 |
|  | 3 | 87,886 | 0.93 | 10,484 |
| CFSAN027350 | 1 | 81,504 | 0.95 | 11,715 |
|  | 2 | 82,543 | 0.96 | 11,641 |
|  | 3 | 87,866 | 1.01 | 11,541 |

To compare assembly performance from Pacbio data, we carried out *de novo* assembly for every strain with each SMRT cell sequenced. Only for CFSAN027350 were the data from three SMRT cells were combined. The overall consensus concordance for the chromosome was between 99.91% and 99.99% and for the plasmids, between 99.95% and 99.99%. All three *de novo* assemblies for CFSAN027346 and CFSAN027350 generated a single contig for the chromosome and the plasmid(s) (Table 6). The three assemblies made using one SMRT cell for strain CFSAN027343 produced a single contig for the plasmid, but 2-3 contigs for the chromosome, due to the presence of a larger repeat (Table 6). To achieve a single contig for the chromosome, we had to combine the three SMRT cells and set the filter for the minimum subread length to 5000 (Supplementary Table 2). Using these settings, the chromosome was able to be closed for CFSAN027343 with a coverage of 400X. After manual closure, a Quiver consensus algorithm was run for each consensus thereby achieving a consensus concordance of 100% with an average coverage from 130X to 170X. It is important to note that the sizes of the chromosome identified for CFSAN027346 and CFSAN027350, and the sizes of the plasmids found after three different SMRT cells runs for each strain were almost identical (variation < 0.0003%).

**Table 6.** Assembly statistics per SMRT cell# for PacBio data using HGAP3.0 and Quiver.

| Strains | Chromosome<br>contig(s) | Plasmids<br>contig(s) | Chromosome<br>size (Mb) | Plasmid(s)<br>size(s) (kb) | Coverage<br>(X) |
| --- | --- | --- | --- | --- | --- |
| **CFSAN027343** |  |  |  |  |  |
| SMRT Cell # 1 | 3 | 1 | 5,351,371;<br>281,039; | 88,847 | 134 |
| SMRT Cell # 2 | 2 | 1 | 81,920<br>5,525,151; | 88,848 | 134 |
| SMRT Cell # 3 | 2 | 1 | 164,081<br>5,525,031;<br>164,076 | 88,847 | 129 |
| **CFSAN027346** |  |  |  |  |  |
| SMRT Cell # 1 | 1 | 2 | 5,592,579 | 1 (96,016), 2 | 147 |
| SMRT Cell # 2 | 1 | 2 | 5,592,570 | (73,152) | 156 |
| SMRT Cell # 3 | 1 | 2 | 5,592,582 | 1 (96,016), 2<br>(73,152)<br>1 (96,017), 2<br>(73,152) | 153 |
| **CFSAN027350** |  |  |  |  |  |
| SMRT Cell # 1 | 1 | 1 | 5,436,071 | 157,534 | 170 |
| SMRT Cell # 2 | 1 | 1 | 5,436,072 | 157,534 | 166 |
| SMRT Cell # 3 | 1 | 1 | 5,436,082 | 157,535 | 176 |

### Detection of Virulence Genes

The output assemblies from our MinION/CANU assembly were used for *in silico* detection of 95 described virulence genes for *E. coli* (39), which include genes reported from all pathotypes of pathogenic *E. coli,* with particular focus on EHECs. To test our hypothesis that MiSeq Nextera XT library sequencing was subpar for obtaining a complete representation of the STEC genomes (chromosome and plasmids), we compared results obtained using MinION assemblies against those obtained from MiSeq assemblies.

The 3 O26:H11 STEC strains were positive for 18 of the 95 virulence genes tested (Table 7). Among those genes were: *astA, cif, eae, ehxA* (plasmid), *espA, espB, espF, espJ, espP* (plasmid), *gad, iha, iss, lpfA, nleA, nleB, nleC, tir,* and *toxB* (plasmid) genes. Other genes were sporadic and strain dependent: *tccP* gene was present in CFSAN027346 and CFSAN027350, *efa1 and katP* (plasmid) genes were only present in CFSAN027343 and CFSAN027346. CFSAN027350 was the only strain found to contain *espI* and *stx2a.* CFSAN027343 and CFSAN027346 carried a different Shiga toxin phage that instead contained the *stx1a* variant.

**Table 7.** Virulence genes present in the O26:H11 STEC MinION assemblies by *in silico* analysis. Plasmid borne genes were: *espP, toxB, katP,* and *exhA.*

| Assemblies<sup>1</sup> | *espI* | *stx2a*<sup>4</sup> | *stx1a*<sup>5</sup> | *tccP* | *efa1* | *katP* |
| --- | --- | --- | --- | --- | --- | --- |
| <sup>2</sup>CFSAN027343a | - | - | + | - | + | + |
| <sup>3</sup>CFSAN027343 | - | - | + | - | + | + |
| <sup>2</sup>CFSAN027346a | - | - | + | + | + | + |
| <sup>2</sup>CFSAN027346b | - | - | + | + | + | + |
| <sup>3</sup>CFSAN027346 | - | - | + | + | + | + |
| <sup>2</sup>CFSAN027350a | + | + | - | + | - | - |
| <sup>2</sup>CFSAN027350b | + | + | - | + | - | - |
| <sup>3</sup>CFSAN027350 | + | + | - | + | - | - |

All assemblies were positive for *astA*, *cif*, *eae*, *ehxA* (plasmid), *espA*, *espB*, *espF*, *espJ*, *espP* (plasmid), *gad*, *iha*, *iss*, *lpfA*, *nleA*, *nleB*, *nleC*, *tir*, and *toxB* (plasmid) genes.
<sup>1</sup>Runs: a) first, and b) second run.
<sup>2</sup>1D Genomic DNA by ligation (SQK-LSK108).
<sup>3</sup>rapid sequencing kit (SQK-RAD002).
<sup>4,5</sup>shiga toxin genes and their variants.

MinION assemblies were congruent and showed the presence/absence of the same genes for each strain (Table 7). We found both nanopore DNA libraries (ligation and rapid) provided sufficient data to assemble and include all virulence genes present, confirming that nanopore technology can provide fast determination of virulence potential by detecting specific virulence genes. Detection of the same virulence genes was also observed with the PacBio assemblies for each strain. However, several virulence genes *(i.e., toxB, tccP, iha,* and *astA)* were not detected in some of the genome assemblies obtained with MiSeq/Nextera XT, pointing to the corresponding library prep as having been responsible for loss of some of these segments.

### *stx* phage identification and location

We assessed whether MinION data could precisely locate where Shiga toxin phages and major pathogenicity islands were located on the genomes, comparing those results with PacBio data – MiSeq was unable to reconstruct individual phages from these strains.

For the MinION-assembled chromosome in CFSAN027343, we detected 20 prophage regions using Phaster (40), of which: 14 regions were intact, 3 regions were incomplete, and 3 regions were questionable (a sample Phaster result is shown in Fig. S3). The *stx* carrying phage was 57.6 Kb located at the *cspG* gene, different from what was previously known for *stx* phage insertions (Table 8). Also, it is interesting to note that although the PacBio sequence gave the same number of prophage regions for this genome, the *stx1* phage was bigger (76.2 kb) and in a different location *(cbpA)* in the resultant PacBio assembly.

**Table 8.**
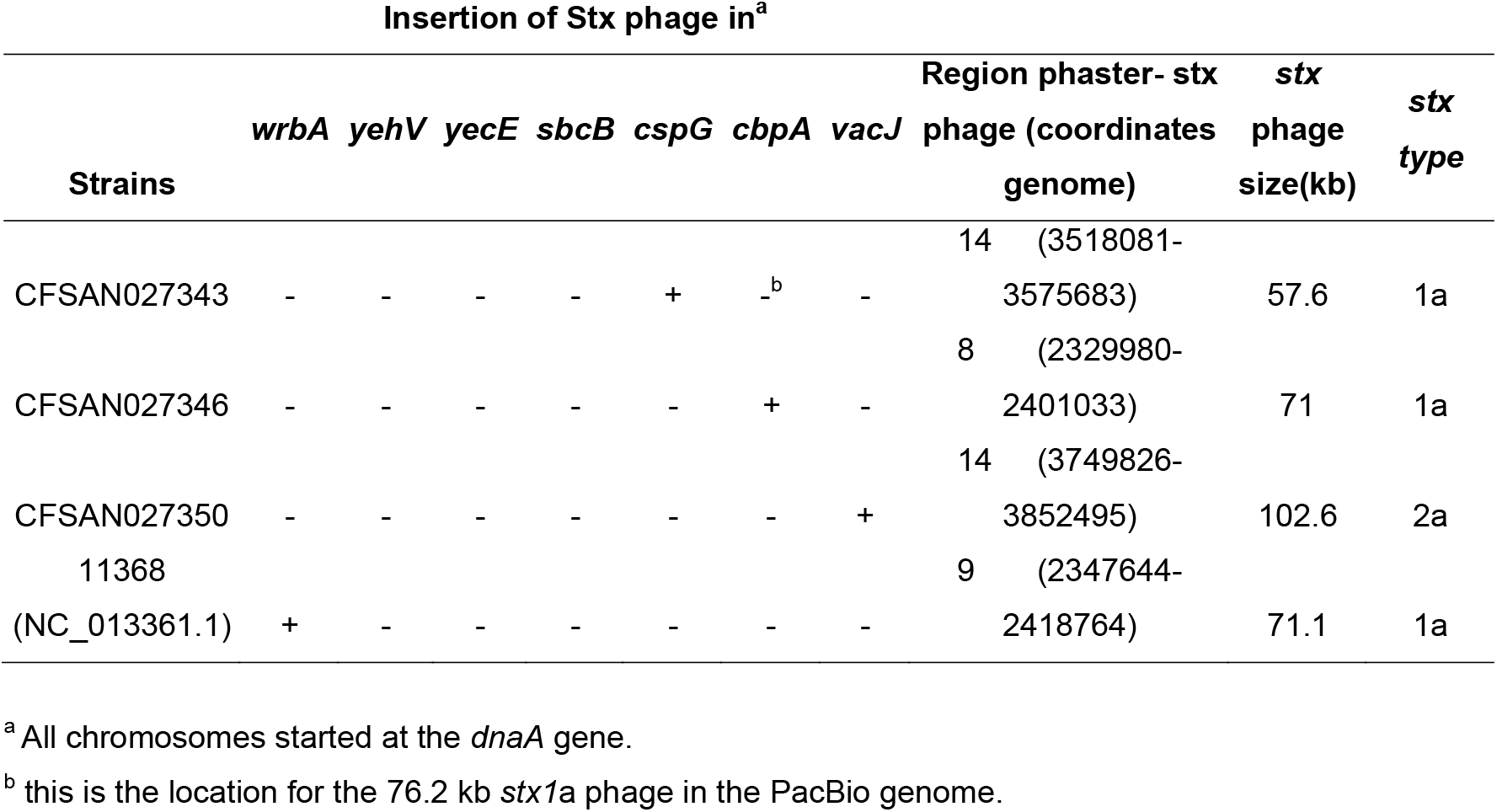
Identification of chromosomal insertion sites for stx phages in the 3 STEC O26:H11 MinION genomes, their *stx* gene type, regions and coordinates in the genome, and their *stx* phage sizes.

| Insertion of Stx phage in<sup>a</sup> |  |  |  |  |  |  |  |  |  |  |
| --- | --- | --- | --- | --- | --- | --- | --- | --- | --- | --- |
| Strains | *wrbA* | *yehV* | *yecE* | *sbcB* | *cspG* | *cbpA* | *vacJ* | Region phaster- stx<br>phage (coordinates<br>genome) | *stx*<br>phage<br>size(kb) | *stx*<br>type |
| CFSAN027343 | - | - | - | - | + | -<sup>b</sup> | - | 14 (3518081-3575683) | 57.6 | 1a |
| CFSAN027346 | - | - | - | - | - | + | - | 8 (2329980-2401033) | 71 | 1a |
| CFSAN027350 | - | - | - | - | - | - | + | 14 (3749826-3852495) | 102.6 | 2a |
| 11368<br>(NC_013361.1) | + | - | - | - | - | - | - | 9 (2347644-2418764) | 71.1 | 1a |

<sup>a</sup> All chromosomes started at the *dnaA* gene.
<sup>b</sup> this is the location for the 76.2 kb *stx1a* phage in the PacBio genome.

In the MinION-assembled chromosome in CFSAN027346, Phaster detected 17 prophage regions (40) : 15 regions were intact, 1 region was incomplete, and 1 region was questionable. PacBio sequence for the same genome was similar, although only 16 prophages were identified (14 intact, 1 incomplete and 1 questionable). Surprisingly, akin to PacBio, that particular *stx* phage was at an unusual insertion site *(cbpA* gene). Nonetheless, both MinION and PacBio found the size of the phage to be 71 kb (Table 8).

In the MinION-assembled chromosome in CFSAN027350, Phaster detected 17 prophage regions (40): 13 regions were intact, 1 region was incomplete, and 3 regions were questionable. PacBio sequence for the same genome identified 16 prophages (11 intact, 5 incomplete; none were questionable). Again, the phage carrying *stx* was located at an unusual insertion site, *vacJ* gene, and this *stx* phage was the biggest among the 3 strains – 102.6 kb (Table 8).

### Detection of antimicrobial genes

*In silico* detection of antimicrobial resistance (AMR) genes using the output assemblies from our MinION pipeline found that only CFSAN027346 carried antimicrobial resistance genes, specifically: aph(3’’)-Ib, aph(6)-Id, blaTEM-1B, *sul2, tetB,* and *dfrA.* All these genes were contained within a smaller plasmid (73 kb) (Table 4 and Table 6) (Fig. S1). Using the assembly generated by the MiSeq/Nextera data, *in silico* analysis showed a similar result but missed the *sul2* gene.

### cgMLST analysis of O26:H11 genomes –MiSeq, Minion, and PacBio

For this section, *de novo* assemblies were produced for the nanopore data using our analysis pipeline described in Figure 1 which includes assembly and Nextera XT polishing of the MinION/CANU assemblies. The phylogenetic relationships among *E. coli* O26:H11/H- strains were determined by a cgMLST analysis shown in Figure 2. This cgMLST was the same as previously published (9). The genome of *E. coli* strain 11368 (NC_013361.1) was used as a reference and has 4,554 genes. The resultant NJ tree showed that the O26 genomes analyzed were highly diverse and polyphyletic and that the assemblies for these 3 strains *(i.e.,* MinION polished assemblies) clustered with the assemblies of the genomes for the same strain generated by the other two technologies that are known to be more accurate *(i.e.,* the MiSeq and PacBio data) (Fig. 2A). However, the assemblies generated by our pipeline were located in longer branches in the same cluster and showed that they still have many errors that differentiated them from their counterparts of high-quality genomes generated by MiSeq or PacBio by 128 – 203 SNPs for CFSAN027343 (Fig. 2C and Fig. S2A) and 92 – 175 SNPs for CFSAN027346 (Fig. 2C and Fig. S2B).

**Figure 1.**
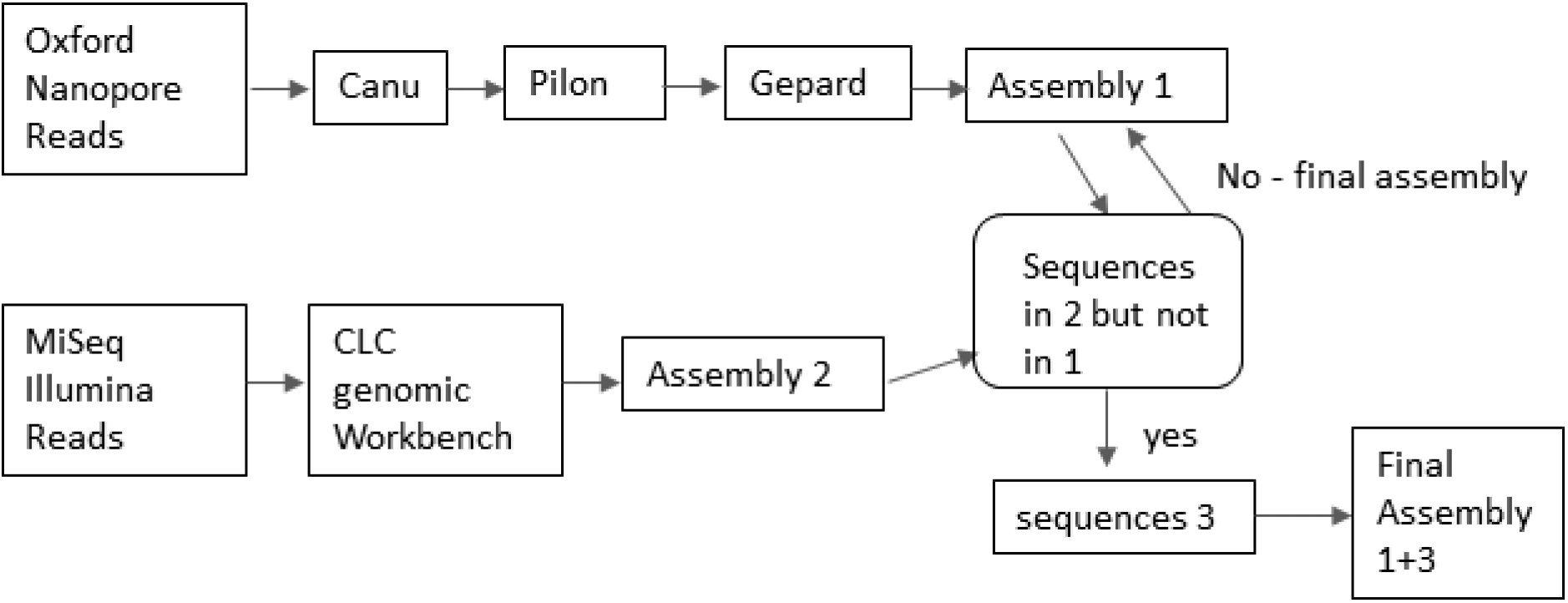
Schematic representation of the analysis pipeline used in this study for assembly and polishing of the MinION sequencing output.

**Figure 2.**
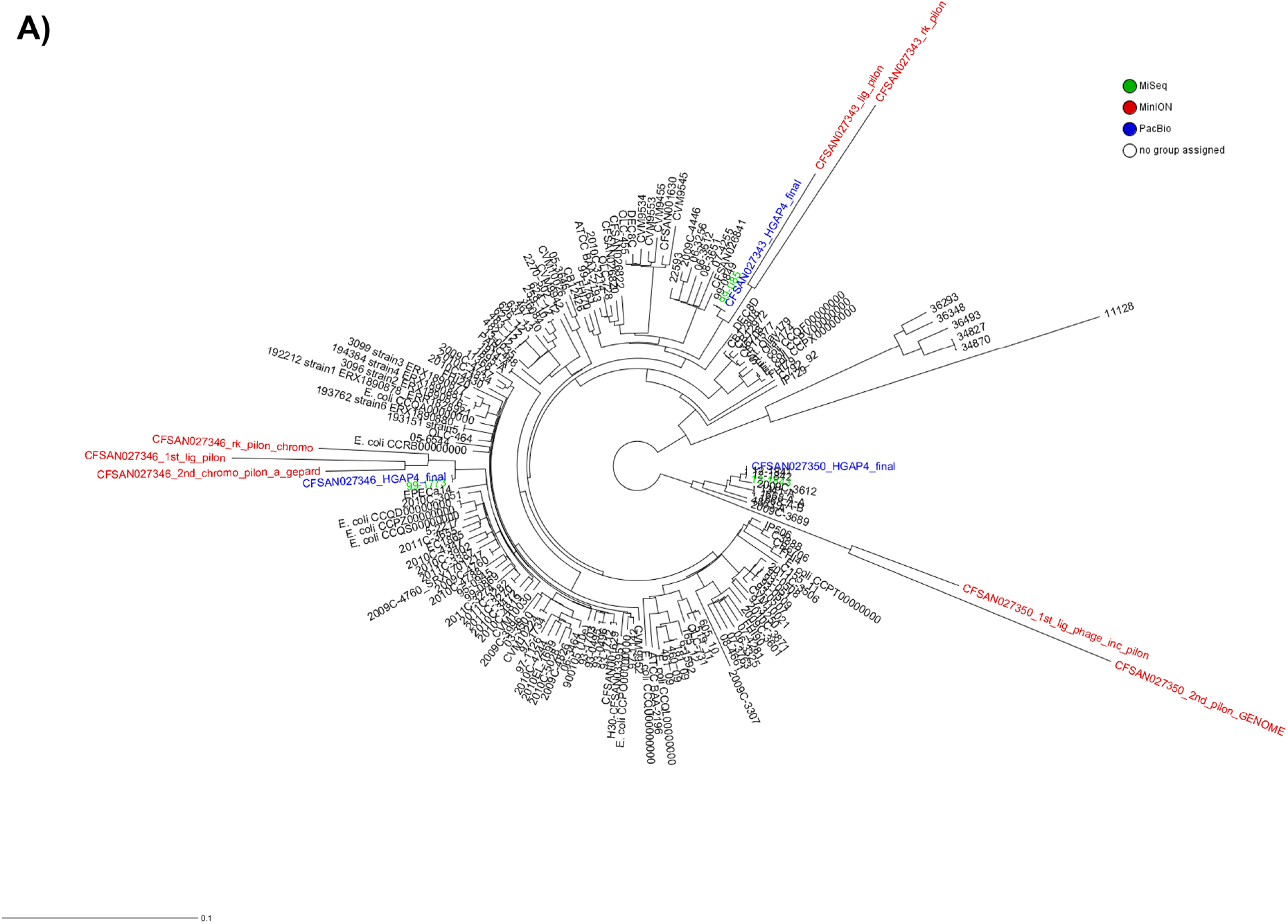
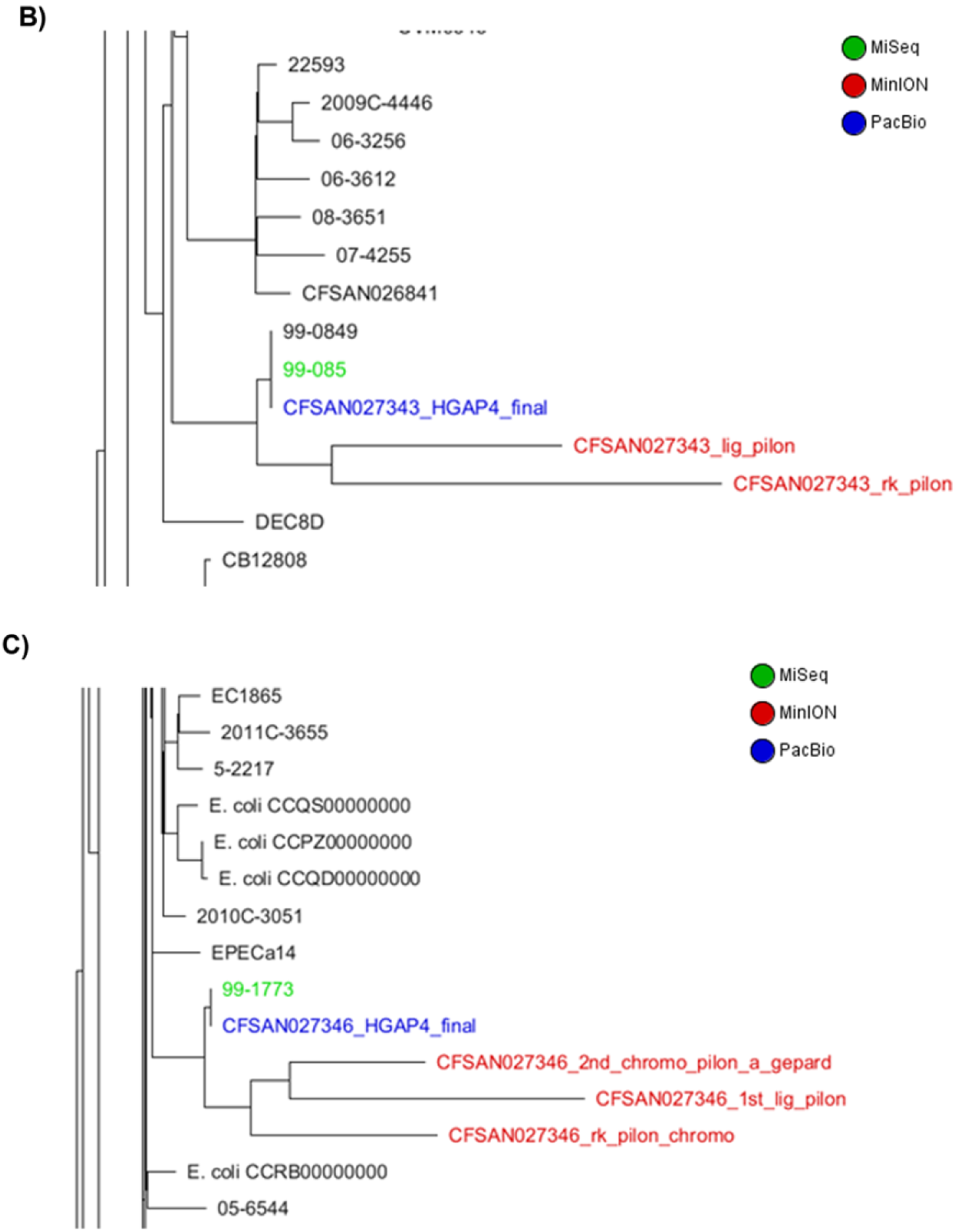
Phylogenetic analysis of the O26:H11/H- *E. coli* strains sequenced in this study by MiSeq, MinION, and PacBio and 195 genomes that are available at GenBank by cgMLST analysis. The evolutionary history was inferred by using the Neighbor-Joining (NJ) tree built using the genetic distance and showing the existence of high diversity and that O26:H11 strains were polyphyletic. A) The genomes generated by any of the 3 technologies still clustered together by the cgMLST analysis. Snapshot of the clusters formed by the genomes generated by the 3 technologies for B) CFSAN27343 and C) CFSAN027346 strains, respectively.

### Synteny comparisons of MinION and PacBio assemblies

A comparison of genome synteny between the assemblies produced by MinION/CANU vs. PacBio was performed using Mauve (41). Overall MinION-generated chromosomes for the 3 strains have almost the same synteny to the ones generated by PacBio (Fig. 3) save for the single exception of CFSAN027343. This strain showed a discordance in chromosomal synteny, with the MinION-generated chromosome appearing more like the Sanger sequence-generated genome strain 11368 (GenBank accession number NC_013361) than the one generated by PacBio (Fig. 3).

**Figure 3.**
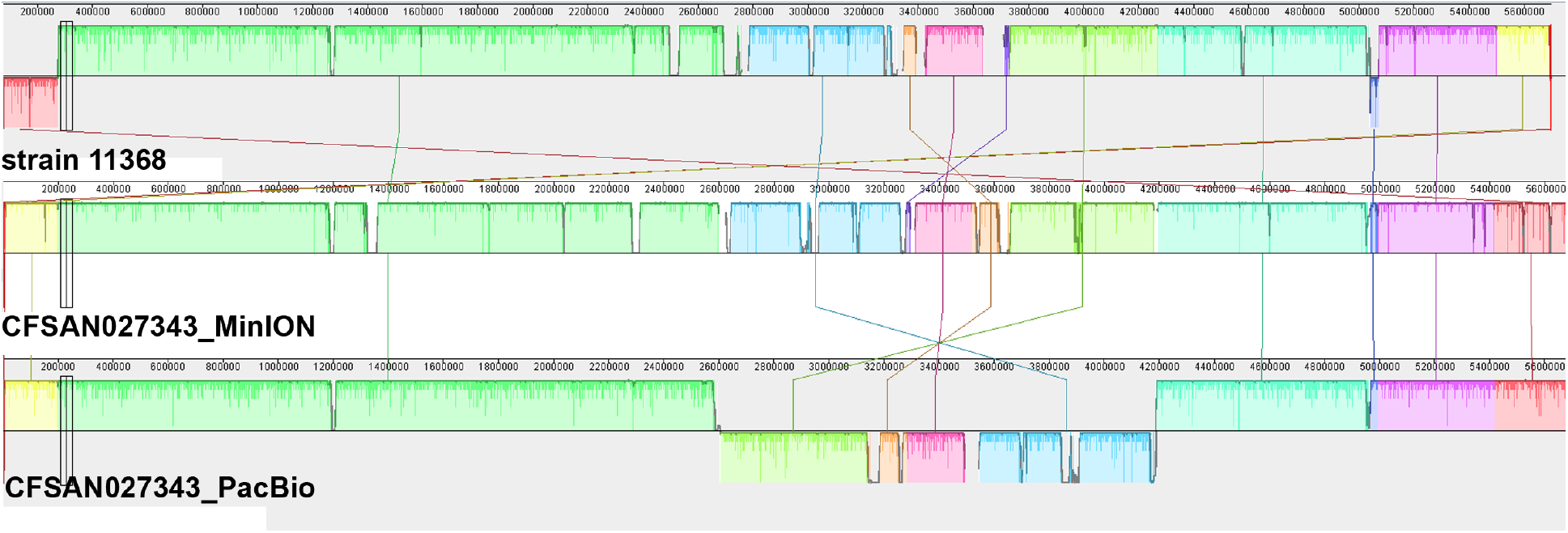
Comparison of the synteny mapping of chromosomes generated by either MinION or PacBio, using the Sanger generated genome for STEC O26:H11 strain 11368 (AP010953) as reference with MAUVE. Each chromosome sequence is laid out in a horizontal track. Matching colors indicate homologous segments and are connected across genomes. Respective scales show the sequence coordinates in base pairs. A colored similarity plot is shown for each genome, the height of which is proportional to the level of sequence identity in that region. Only strain CFSAN027343 synteny is shown for illustration purposes (The other two strains (CFSAN027346 and CFSAN027350) synteny can be found in Fig. S4).

### Novel plasmid in O26:H11 (155 kb)

The virulence plasmids of CFSAN027343 and CFSAN027346 were 88 kb and 96 kb, respectively. Interestingly, the virulence plasmid for CFSAN027350 was much larger (157 kb). Both pCFSAN027343 and pCFSAN027346, however, carried these 4 virulence genes: *exhA, espP, toxB,* and *katP,* while pCFSAN027350 carried only *exhA, espP,* and *toxB.* Comparative analyses of the 3 plasmids showed that pCFSAN027350 was very different from the other 2 plasmids (Fig. 4), possessing extensive unique regions. Based on these data, pCFSAN027350 contains 214 annotated ORFs with multiple transposons and insertion elements and has a G + C content of 48.4 % (Table 9).

**Figure 4.**
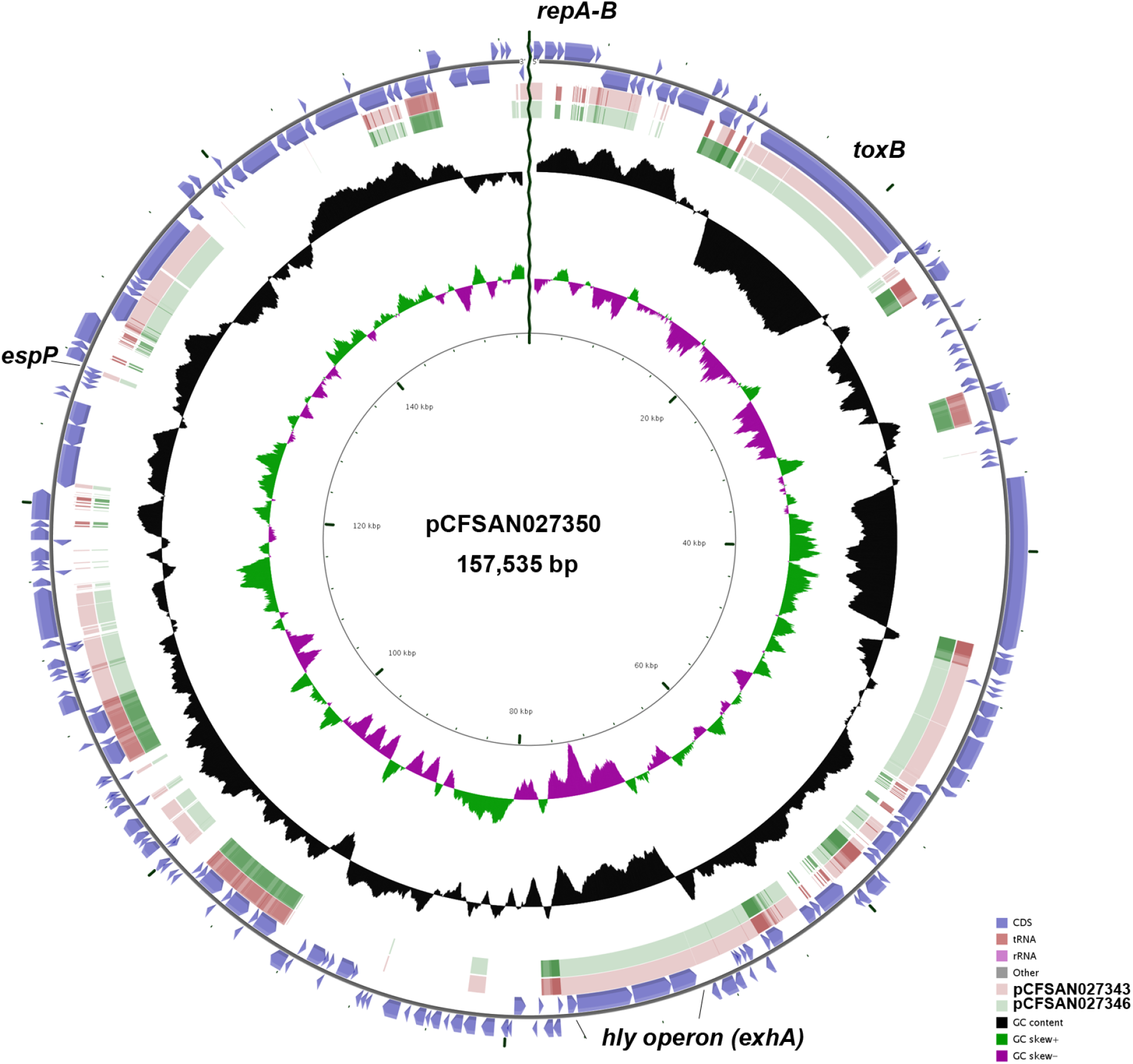
Circular map of virulence plasmid pCFSAN027350 compared to the other two virulence plasmids (pCFSAN027343 and pCFSAN027346), generated using CGView (66). Blue block arrows in the outer circle denote coding regions in the plasmid, indicating the ORF transcription direction. G+C content is shown in the middle circle and the deviation from average G+C content (47.71%) is displayed in the innermost circle. BLAST comparisons with the other two EHEC plasmids are shown in light red (pCFSAN027343) and green (pCFSAN027346).

**Table 9.**
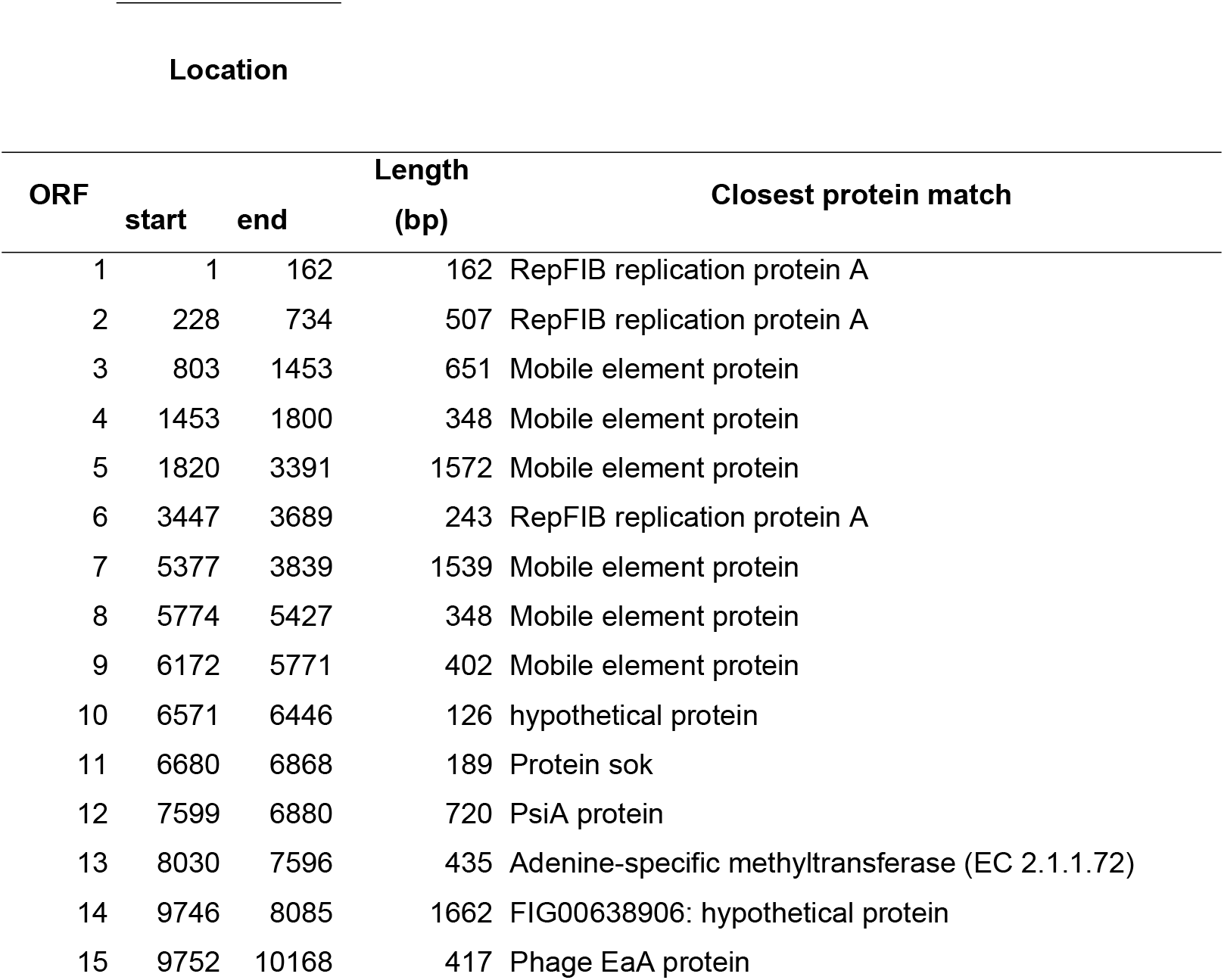
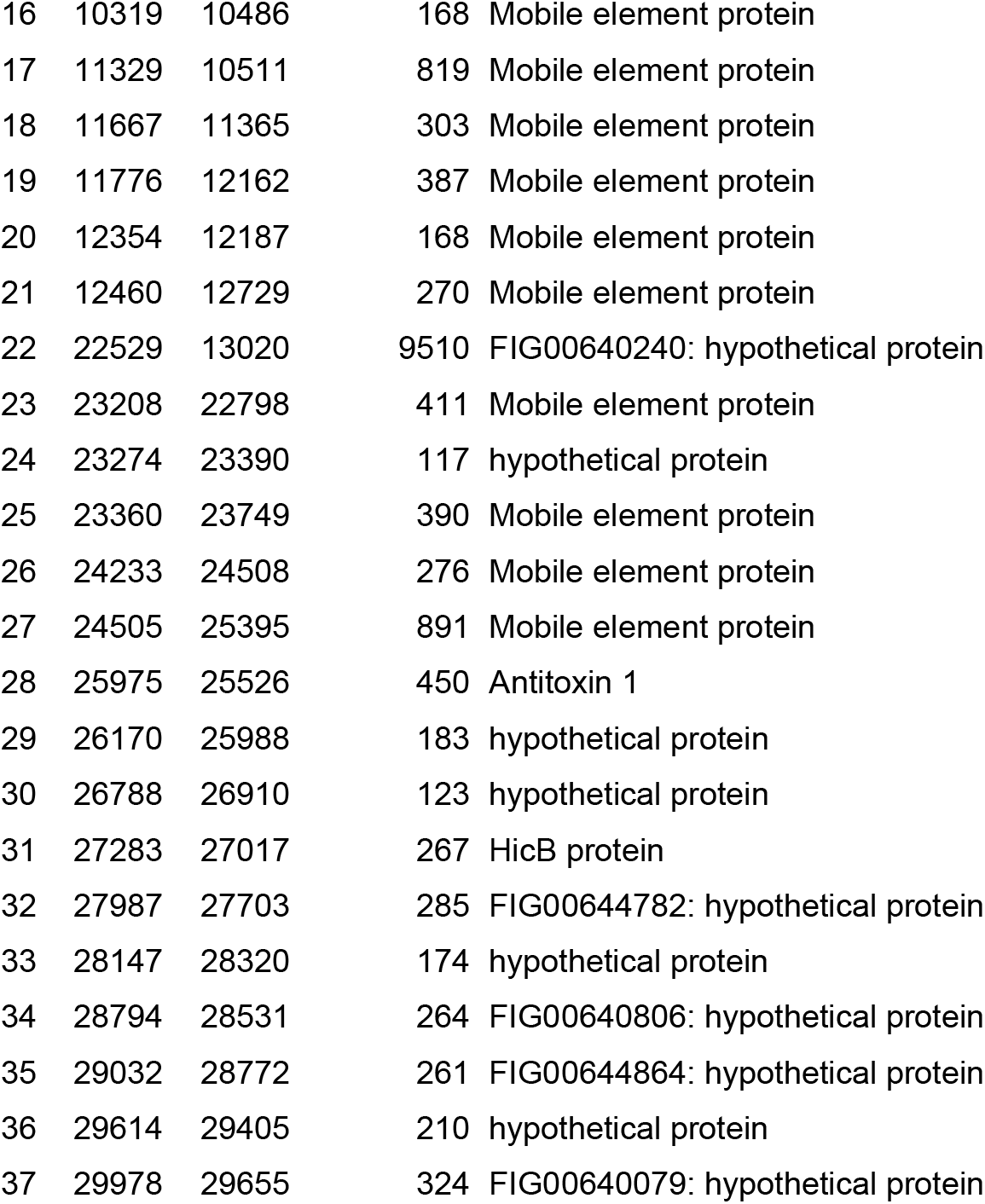
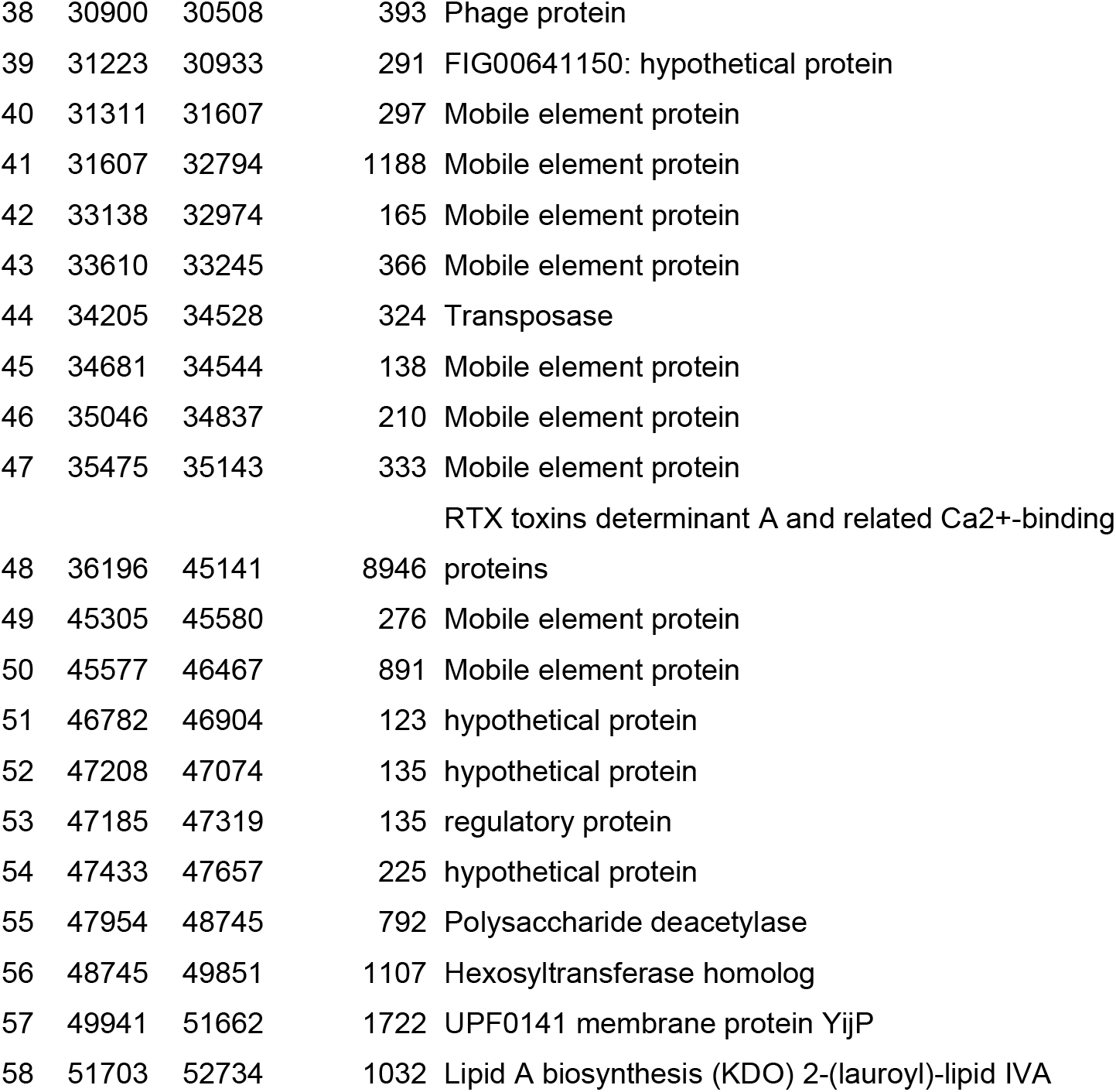
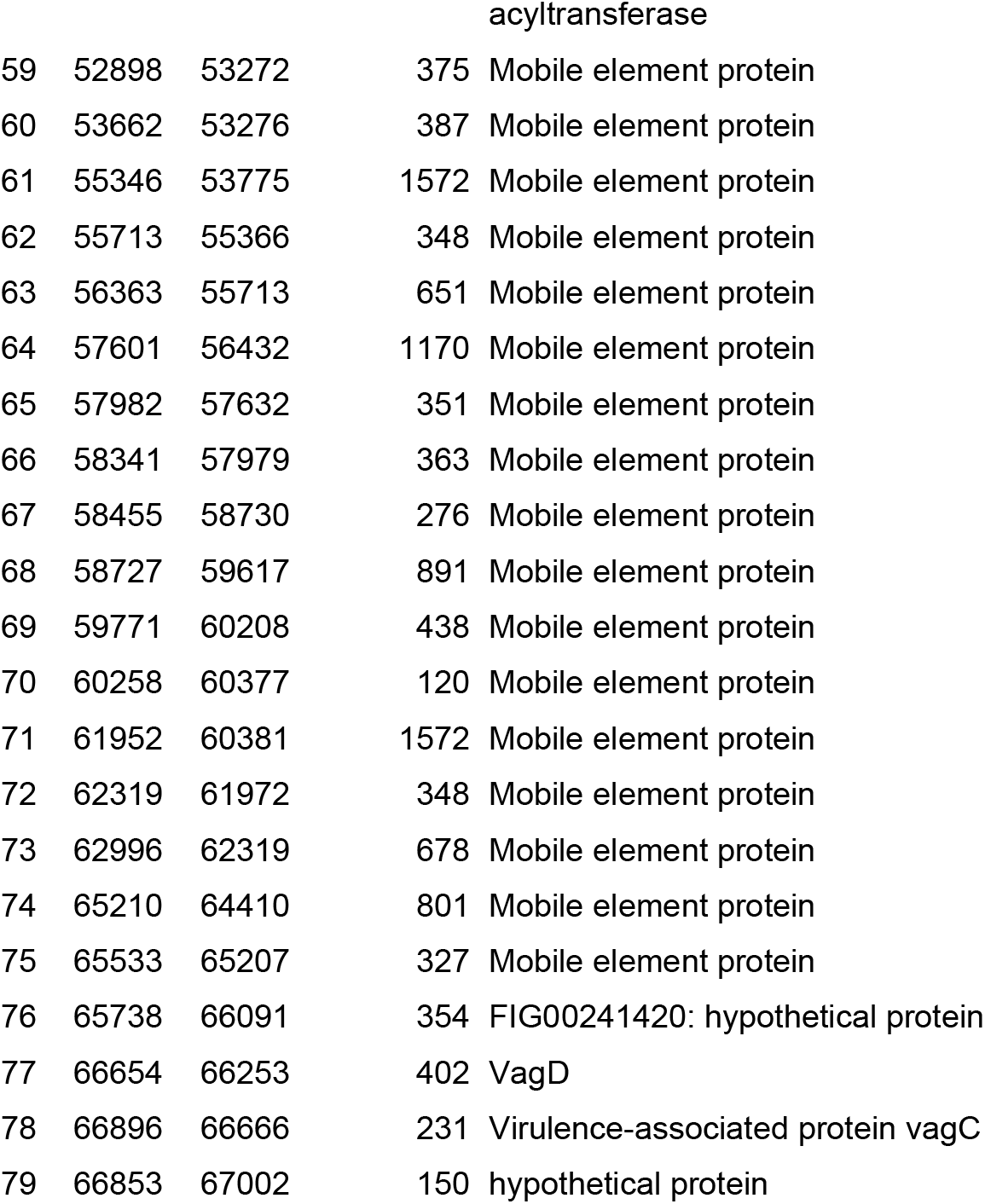
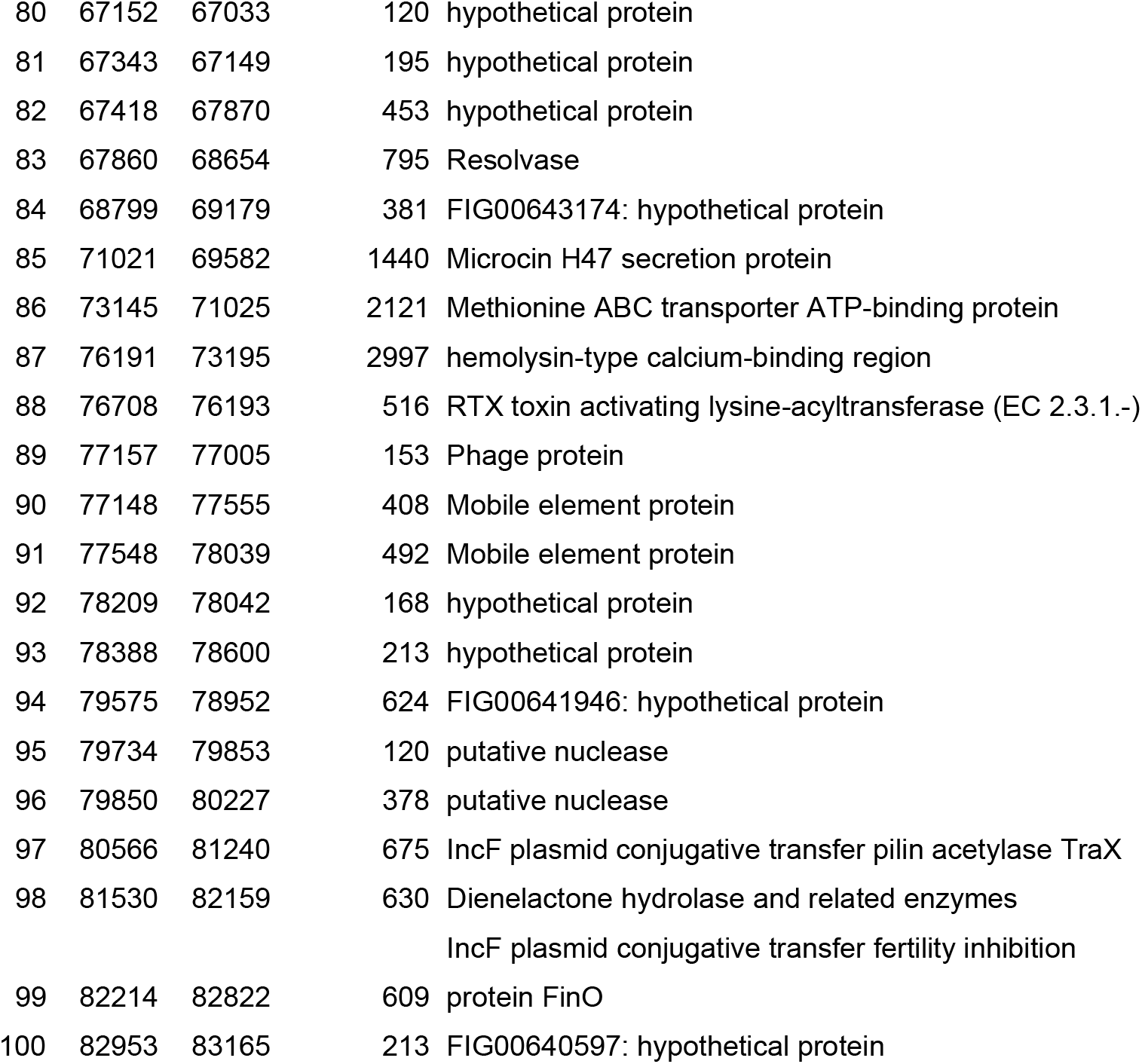
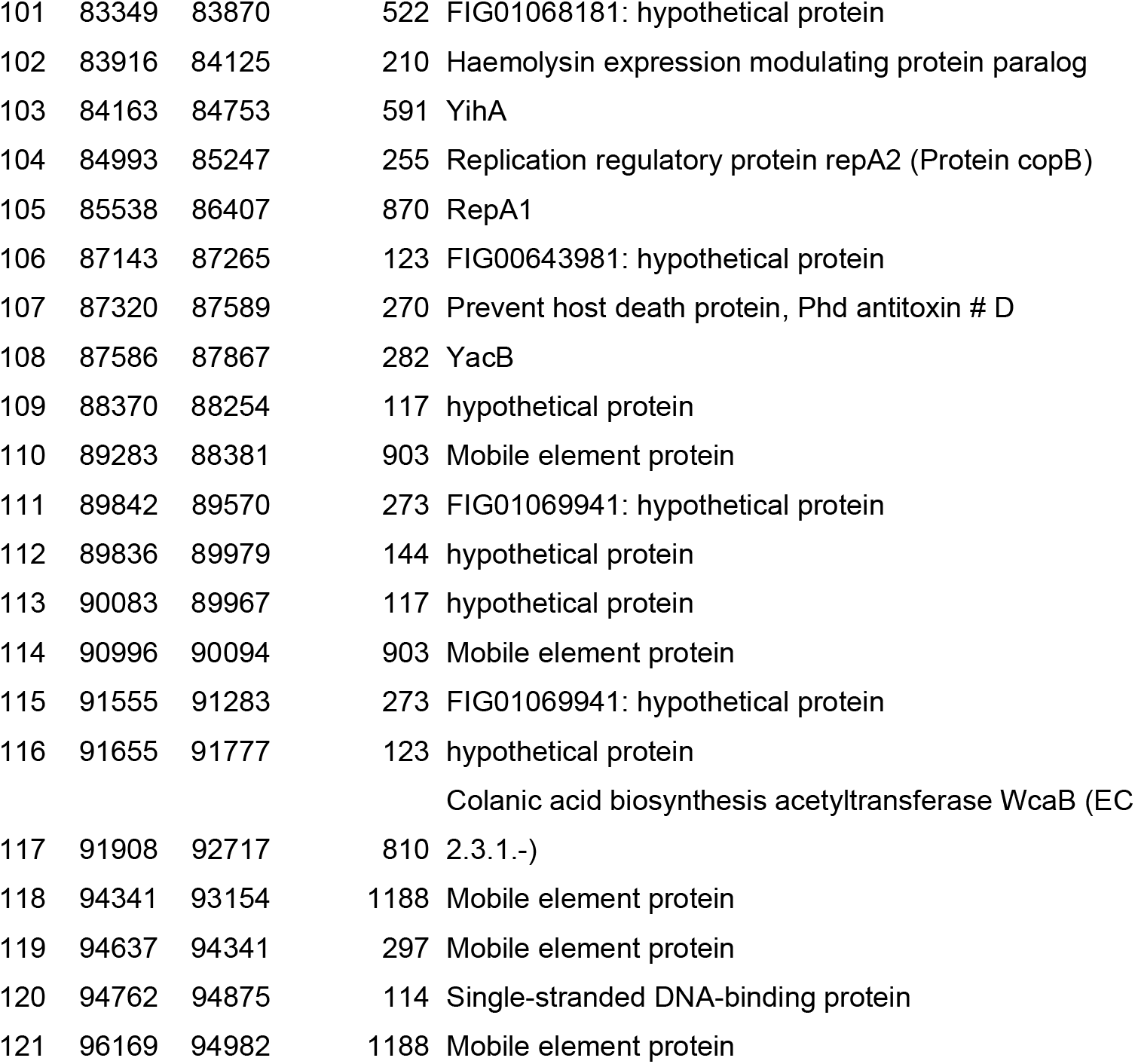
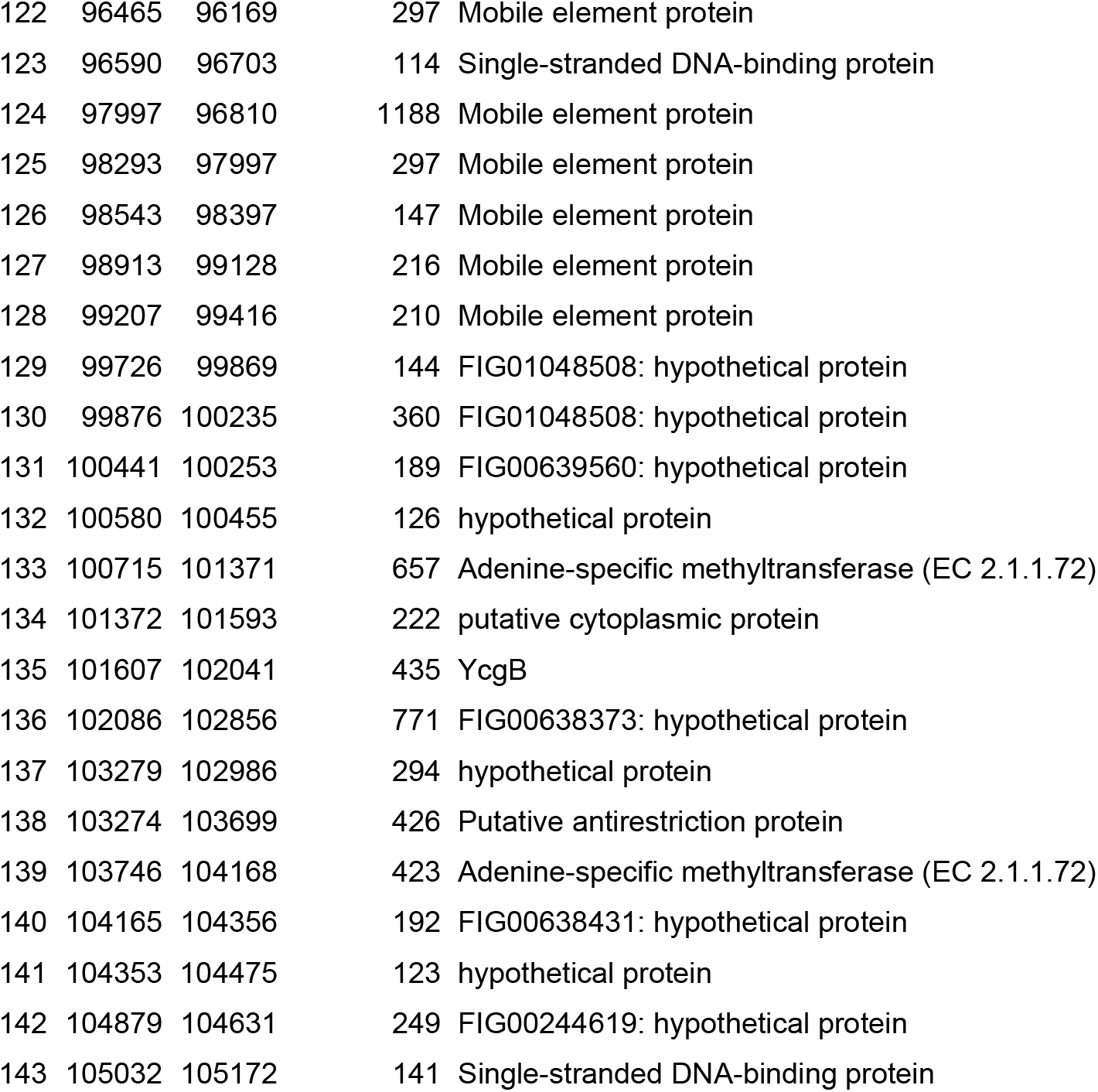
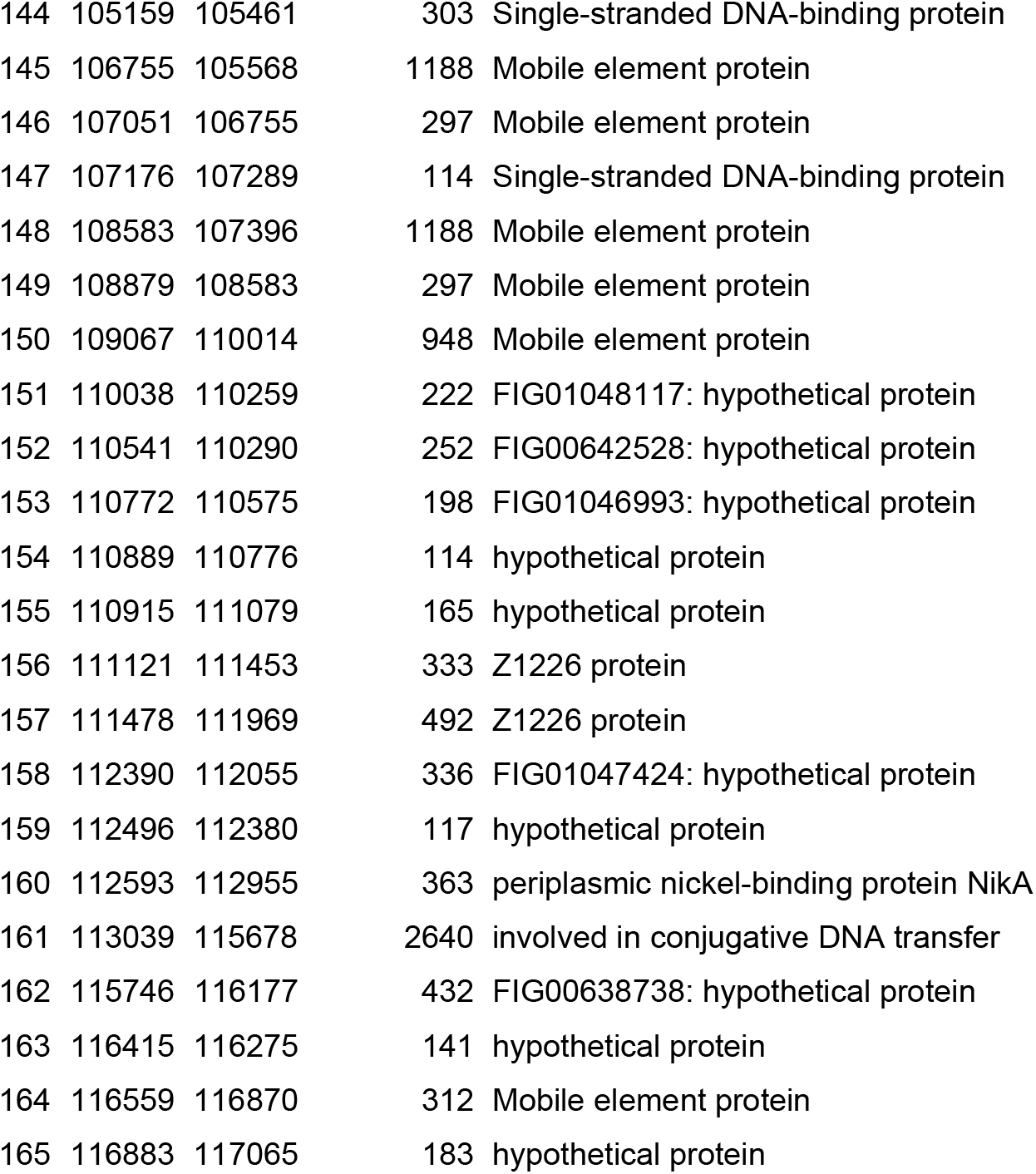
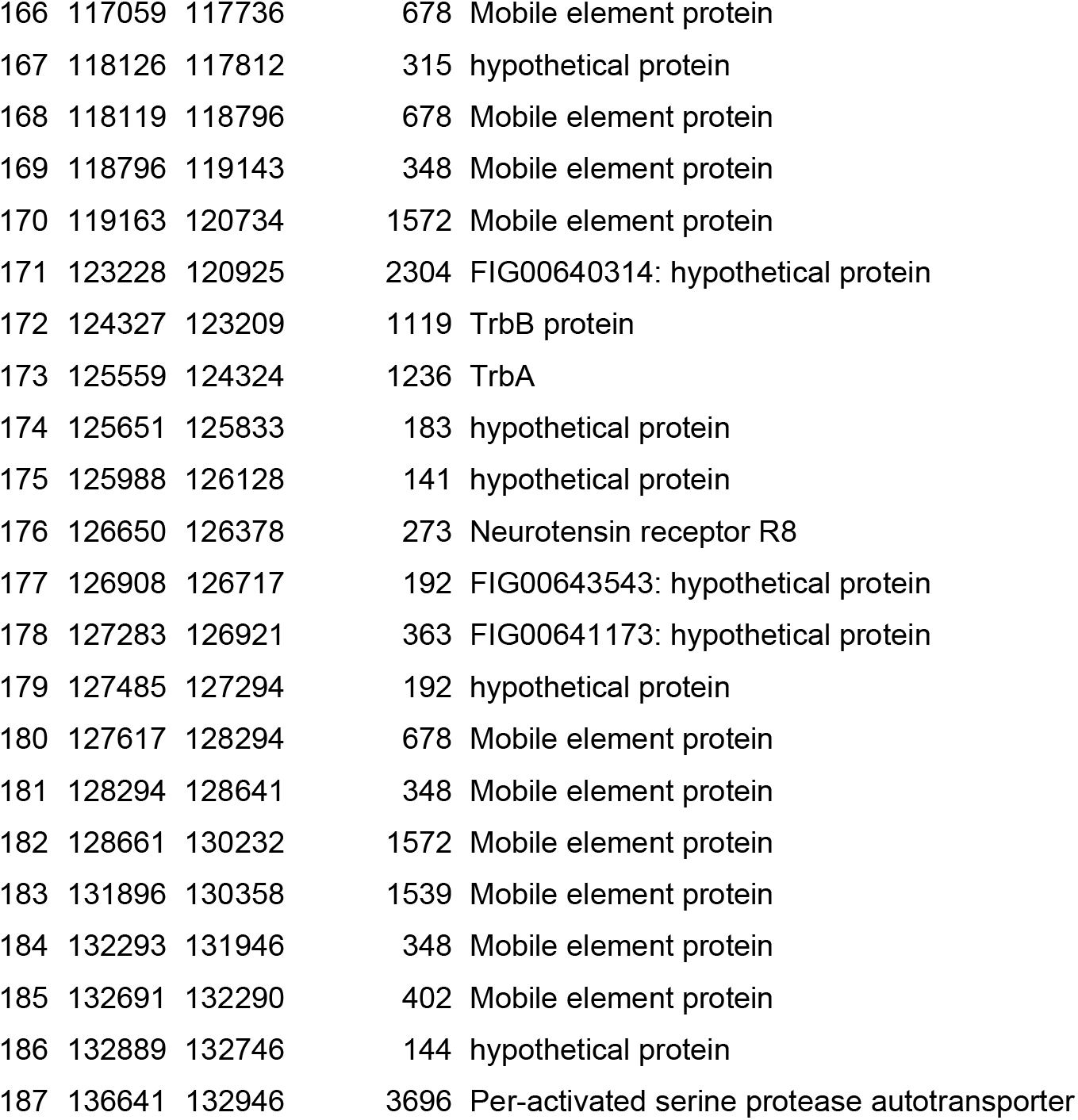
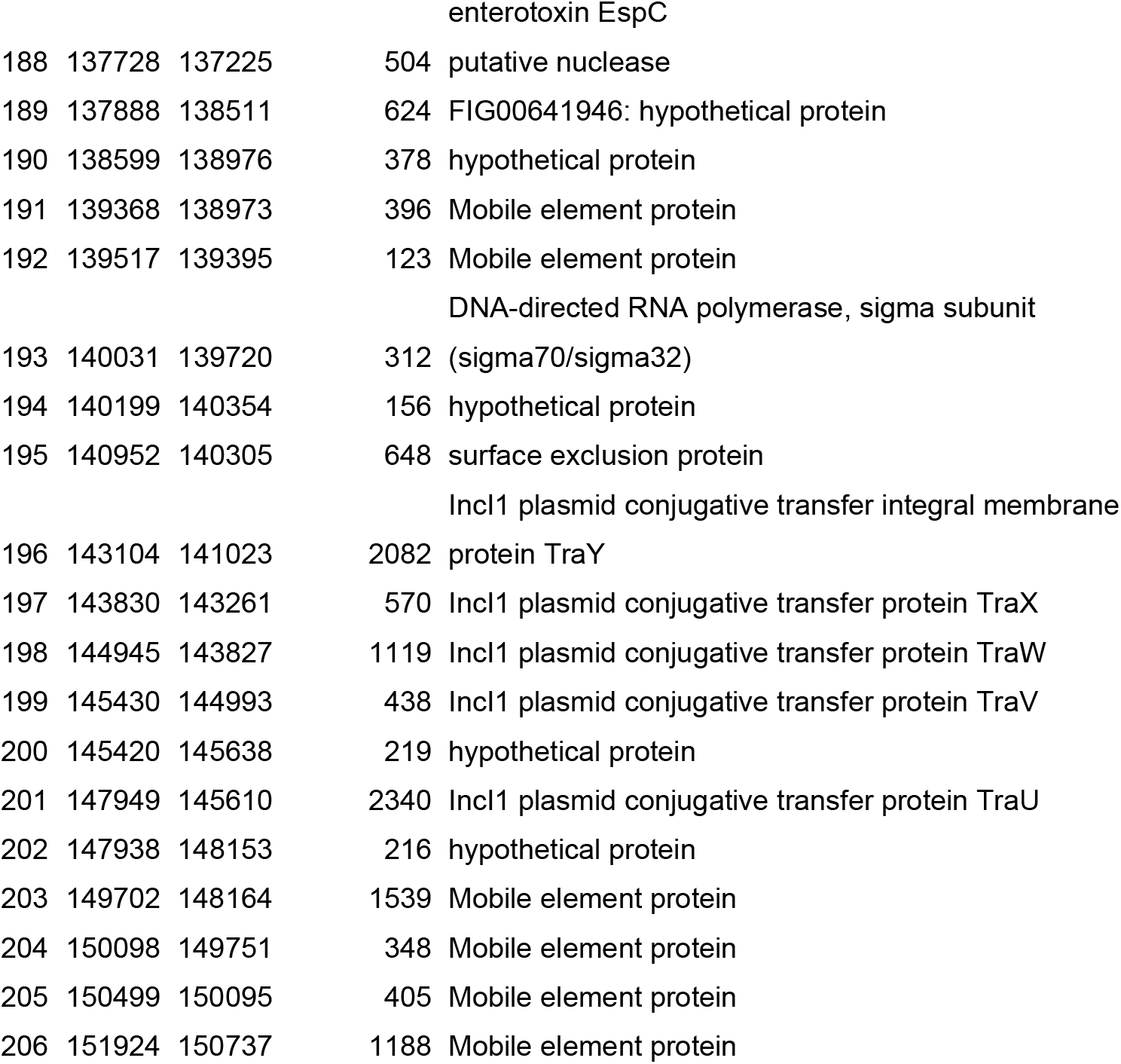
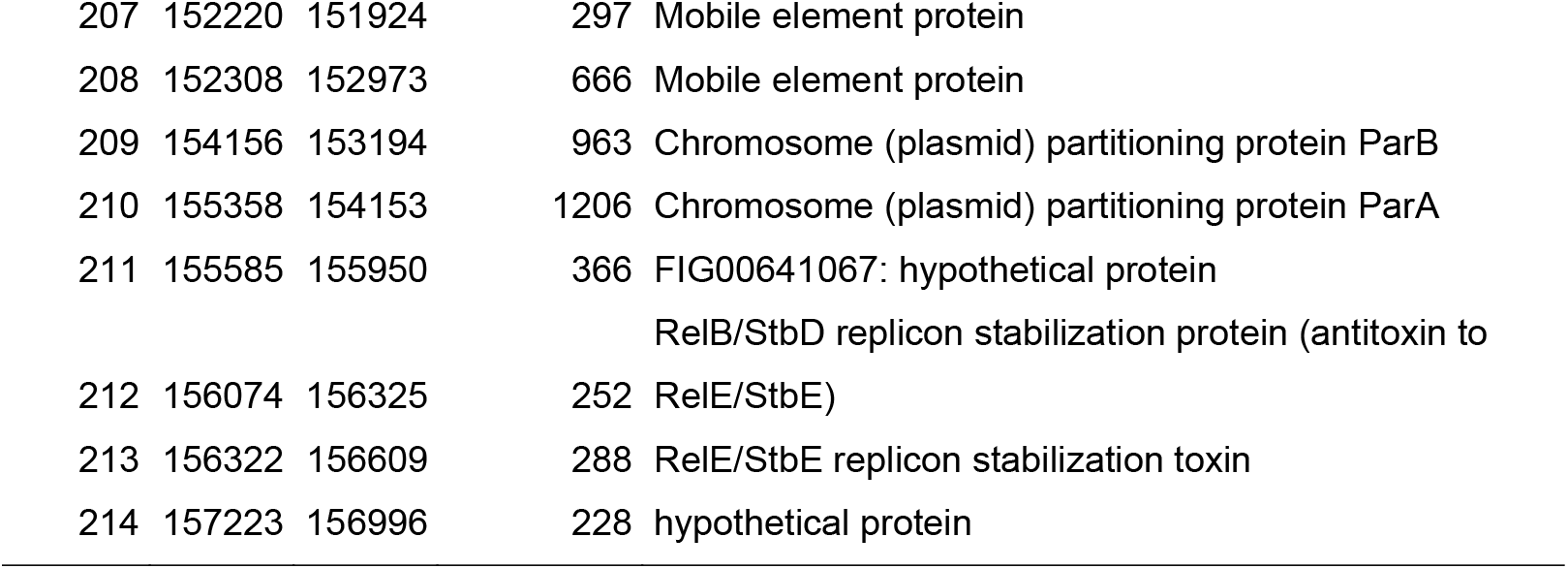
Summary of ORFs in pCFSAN027350 as annotated by RAST^59^.

## DISCUSSION

Based on MLST analyses, strains of STEC O26:H11/H- can be separated into several STs; of these, ST21 and ST29 have been associated with disease in humans (7). Therefore, it is especially important to be able to rapidly characterize all detected STECs O26:H11/- (both chromosome and plasmid(s), if present) to determine whether those clones or strains circulating in particular locations are likely to be threats to human health. Here, we tested the use of such a technique *(i.e.,* Nanopore sequencing) to obtain the complete genomes of 3 unrelated STEC O26:H11 strains followed by subsequent comparisons to those obtained by PacBio and MiSeq WGS platforms. The 3 strains belonged to the EHEC O26 group in accordance with the O26:H11/- cgMLST scheme reported previously by Gonzalez-Escalona et al. in 2016 (9).

We conducted Nanopore testing using a MinION device connected to a laptop using two different DNA sequencing kits (the 1D ligation kit and the rapid sequencing kit RAD002) in the study. Resultant genomes were able to be closed using any of the DNA sequencing kit deployed here save for CFSAN027350 which failed to generate a complete genome in a single contig for the chromosome – was contained in 15 contigs using the rapid kit (RAD002). Nonetheless, the virulence plasmid for that strain/run was contained in a single contig. One explanation for this discrepancy could be that the WGS output was very low at <0.14 Gb for reads above 5,000 bp *(i.e.,* 21,759 total reads). Also, the minimum amount of reads that were necessary for closing the STEC genome was around 58,000 reads above 5,000 bp for the RAD002 kit. The number of reads generated by the 1D ligation kit was around 5 times higher than that for the RAD002 kit, which allowed us to estimate that, in principal, we could run up to 5 different strains per flow cell thus reducing the cost per genome to ∼200 USD. This assumption is reinforced by the newer released RAD004 kit, which produces even higher and better-quality output that its predecessor (unpublished observations).

In the last 4 years, Pacbio sequencing, combined with *de novo* assembly, has been an attractive method for closing bacterial chromosomes and the plasmid(s) that they may carry. Indeed, in this study, we verified that by using PacBio technology with P6-C4 chemistry and *de novo* assembly, the three *E. coli* genomes including their plasmids studied here were able to be completely sequenced and closed. The three assemblies obtained for each isolate using the raw sequence data from each SMRT cell run, respectively, showed high consistency and reproducibility with a coverage of above 100X and an average read length of 10 to 12 kb. Moreover, all chromosomes and plasmids could be completely closed with the sequence data from one SMRT cell except for the single chromosomal sequence run from CFSAN027343. Fragmented assemblies can occur when the repetitive region in some bacteria is longer than the read lengths. Pathogenic *E. coli,* in fact, are known for carrying repetitive regions within the chromosome and read lengths obtained from the PacBio sequencer, at times, are not long enough to overcome this challenge. Albeit, Pacbio is a stand alone instrument that does not require finishing or polishing with additional WGS data from another source.

Using MinION-generated genomes, polished with Nextera XT illumina reads, allowed us to accurately and efficiently determine the virulence genes present in each strain, as well as which genes were located on plasmid(s) or on the chromosome (see Table 7). We also were able to determine the presence of AMR in one of the strains (CFSAN027346) and that was located on an extra plasmid carried solely by that strain (Fig. S1). The other two STEC strains carried a single virulence plasmid albeit with different sizes and gene compositions. It was also noteworthy that none of these strains carried the *etpD* gene (coding for a Type II secretion protein) in the plasmid, indicative of the presence of the European clone (9,35,45,46). From the nanopore data analyzed here, the virulence profile for each strain was determined as follows: CFSAN027343 (ST21, stx^1a^+, *eae-beta1+,* plasmid gene profile *ehxA+, espP+, katP,* and *toxB+*), CFSAN027346 (ST21, stx^1a^+, *eae-beta1+,* plasmid gene profile *ehxA+, espP+, katP,* and *toxB+),* and CFSAN027350 (ST29, stx^2a^+, *eae-beta1+,* plasmid gene profile *ehxA+, espP+,* and toxB+).

Besides differences in their Shiga toxin phage content, we detected other virulence genes unique to some of the 3 STECs analyzed here. Among them was the *tccP* gene, coding for an effector protein (Tir-cytoskeleton coupling protein), found commonly in O157:H7 EHEC strains (47,48) and which was present in strains CFSAN027346 and CFSAN027350. The *tccP* gene plays a direct role during EHEC infection inducing actin polymerization by coupling Tir to the actin cytoskeleton (49). The presence of this gene, together with *espJ, stx1a* or 2a, intimin, and Tir, makes these strains highly pathogenic. Another important illustration of the mosaic of virulence factors found within STEC O26:H11 strains is that two important genes for intestinal colonization *(efa1/lifA -* a protein with putative glycosyltransferase activity that has an important role in intestinal colonization), *and katP- (a* catalase-peroxidase, which might help EHEC O157:H7 to colonize host intestines by reducing oxidative stress), were present in both CFSAN027343 and CFSAN027346 (50,51).

We also sequenced the same 3 strains using PacBio, which is considered the gold standard for closing bacterial genomes (10,11,52). These genomes were compared to the ones generated by nanopore sequencing. Overall, there was fair agreement with the sequences in both the chromosome(s) and plasmid(s) generated from the 3 STEC strains. This was evidenced by the extremely high synteny as well as virulence genes profiles, AMR genes, and overall plasmid content. Despite this extraordinary congruence, some discrepancies were observed for CFSAN027343 where there was a fragment (∼ 1.5 Mb) that appears to be reversed in direction on the chromosome between the two, influencing the resultant calling of the *stx* phage and with MinION stx-phage being smaller (57.6 kb) than the PacBio stx-phage (75.6 kb).

As observed by other authors, sequencing accuracy continued to be an issue for the genomes generated by nanopore sequencing here. That is, PacBio maintained 99% accuracy (1 SNP observed by cgMLST) while MinION genomes revealed expectedly lower accuracy (92 – 203 SNPs by cgMLST) (Fig. S2). Without Illumina Nextera XT polishing, we are unable to accurately place the MinION assemblies -using this cgMLST scheme- into the phylogenetic tree as clustering relies critically on sequence calling using complete ORFs. One of the main drawbacks of MinION nanopore sequencing is that the assemblies contained numerous artifactual indels that will readily translate into incorrect allele calls. This will, in turn, affect the correct clustering of those isolates in the phylogenetic tree. Continued improvements in nanopore-based sequencing chemistry *(e.g.* RAD004) as well as development of alternative base-calling algorithms such as Scrappie – an open-source transducer neural network (53) may further mitigate this problem. This latter development is designed to aid in the correction of longer homopolymeric runs, one of the main challenges of the nanopore platform. On the other hand, it is equally important to note that PacBio also has continued to mitigate caveats including to time of labor, and read length with recent advances in Sequel chemistry (SMRTbell Express Template Prep Kit, Sequel Sequencing and Binding Kit 3.0 and SMRT Cell 1M v3).

By using MinION sequencing we were able to easily identify the location and composition of the shiga toxin carrying phages in the STEC O26:H11 strains. Contrary to what has been observed for EHEC O157:H7 and non-O157:H7 STECs, the location on the chromosome of the stx phages (54) were in different regions of the chromomere for each strain in this study. According to Bananno et al (2015), nine Stx phage insertion sites have been described in STECs strains genes: *wrbA, yehV, yecE, sbcB, Z2577, ssrA, prfC, argW, torS-torT* intergenic region. For the prototypic Sanger sequence STEC O26:H11 strain 11368, the Stx phage is located at the *wrbA* gene, a very common observed site for *stx* phage insertions in STECs (54). The Stx phage in these 3 strains (sequenced in this study) were located in novel locations on the chromosome *(cspG* – CFSAN027343, *cbpA*- CFSAN027346, and *vacJ* – CFSAN027350), highlighting once more the high diversity among STECs O26:H11/- strains.

Finally, with both MinION and PacBio, we were able to identify a large virulence plasmid in strain CFSAN027350 (∼ 157 kb) (Fig. 4). The presence of a large virulence plasmid in O26:H11 strains is not uncommon as was observed for strain H30 (pO26-Vir – 168 kb) (36), which contained 5 additional plasmids. As was observed with pO26-Vir, pCFSAN027350 showed a mosaic structure, with many fragments of the plasmid matching other plasmids available at GenBank and contained multiple transposons and insertion sequences, evidence of the mobility of some of the regions (e.g. *toxB* gene was surrounded by IS elements at both ends of the gene). Nevertheless, this plasmid was highly similar (99%) to another plasmid reported for a clinical strain of O26:H11, albeit of larger size (∼181 kb, strain 2013C-3277 plasmid unnamed3, CP027334.1).

Overall, the high degree of correlation we found between these two long-read methods for determining plasmids, virulome, antimicrobial resistance genes, and phage composition in STEC O26 strongly indicates that the MinION sequencing technology is an excellent solution for rapidly determining STEC O26 closed genomes and performing comprehensive analysis of their genomic markers. The MinION devices are robust enough to be used to monitor viruses in remote areas (16), yet can provide most of the applications provided on the PacBio platform (methylation patterns, metagenomics, finding location of pathogenicity islands, resolving plasmids, among other applications) (13,17,23–25,28). To be able to rapidly provide an assessment of possible virulence, antimicrobial resistance potential and disease risk posed by any STEC using this nanopore sequencing technology, is of paramount importance for the protection of the public health and for the tracking of existent and new clones in any geographical region.

## ACKNOWLEDGMENTS

The study was supported by funding from the MCMi Challenge Grants Program Proposal #2018-646 and the FDA Foods Program Intramural Funds. The authors thank Dr. Lili Fox Vélez for her scientific writing assistance on this manuscript. The views expressed in this article are those of the authors and do not necessarily reflect the official policy of the Department of Health and Human Services, the U.S. Food and Drug Administration (FDA), or the U.S. Government. Reference to any commercial materials, equipment, or process does not in any way constitute approval, endorsement, or recommendation by the FDA.

## MATERIALS AND METHODS

### Bacterial strains and media

*E. coli* strains CFSAN027343, CFSAN027346, and CFSAN027350 (Table 1) were purchased from the *E. coli* Reference Center (Pennsylvania State University, University Park, PA). These strains were revived in Tryptic Soy Broth and grown overnight at 37°C.

### DNA preparation

Genomic DNA from each strain was isolated from overnight cultures using the DNeasy Blood and Tissue Kit (Qiagen, Valencia, CA), following the manufacturer’s instructions. The extracted DNA was stored at −20 °C until used as a template for whole genome sequencing. The concentration was determined using a Qubit double-stranded DNA HS assay kit and a Qubit 2.0 fluorometer (Thermo Scientific), according to manufacturer’s instructions.

### MinION nanopore whole genome sequencing and contigs assembly

The genomes of the strains were sequenced using a MinION device (Nanopore, Oxford, UK), with FLO-MIN106 (R9.4) flow cells, according to the manufacturer’s instructions, at > 50 X – average coverage. The sequencing libraries were prepared using either the 1D Genomic DNA by ligation kit (SQK-LSK108) library chemistry or the rapid sequencing kit (SQK-RAD002) according to the manufacturer’s instructions. An exception was for the 1D ligation kit where we omitted the shearing step and the initial step was the End-prep step, since the DNA extraction step already sheared the DNA. The DNA input was 1 ug per DNA library for all 1D Genomic DNA by ligation kit (SQK-LSK108) and 0.4 ug per rapid sequencing kit (SQK-RAD002) library. Each prepared library was loaded into a FLO-MIN106 flowcell (R9.4) for a 48-hour run. *E. coli* strains CFSAN027346 and CFSAN027350 were sequenced in duplicate using the ligation kit, and a single sequencing of each strain was conducted suing the RAD002 kit (Table 2). The base calling was performed using Albacore software (Nanopore, Oxford, UK). The fastq files were generated from the base called sequencing fast5 reads using Poretools (55). Genomic sequence contigs were *de novo* assembled using default settings within the CANU program (56) v1.6. Determination of the overlapping regions of the chromosome and plasmids were carried out using Gepard (57). The resultant assemblies from CANU were corrected for errors using Pilon (58) and the MiSeq data generated from those strains. Our de novo assembly and polishing pipeline are shown in Figure 1.

### PacBio whole genome sequencing and contigs assembly

The strains were sequenced on the Pacific Biosciences (PacBio) *RS* II Sequencer, as previously described (2,11,52). Specifically, we prepared the library using 10 μg genomic DNA that was sheared to a size of 20kb fragments by g-tubes (Covaris, Inc., Woburn, MA) according to the manufacturer’s instruction. The SMRTbell 20-kb template library was constructed using DNA Template Prep Kit 1.0 with the 20-kb insert library protocol (Pacific Biosciences; Menlo Park, CA, USA). Size selection was performed with BluePippin (Sage Science, Beverly, MA). The library was sequenced using the P6/C4 chemistry on 3 single-molecule real-time (SMRT) cells with a 240-min collection protocol along with stage start.

Analysis of the sequence reads was implemented using SMRT Analysis 2.3.0. The best de novo assembly was established with the PacBio Hierarchical Genome Assembly Process (HGAP3.0) program using the continuous-long-reads from the four SMRT cells. The assemblies outputs from HGAP contains overlapping regions at the end which can be identified using dot plots in Gepard (57). Genomes were checked manually for even sequencing coverage. Afterwards the improved consensus sequence was uploaded in SMRT Analysis 2.3.0. to determine the final consensus and accuracy scores using Quiver consensus algorithm (52).

### MiSeq whole genome sequencing, contig assembly and annotation

The genomes of the strains were sequenced using an Illumina MiSeq sequencer (Illumina, San Diego, CA), with the 2×250 bp pair-end chemistry according to manufacturer’s instructions, at approximately 80X average coverage. The genome libraries were constructed using the Nextera XT DNA sample prep kit (Illumina). Genomic sequence contigs were *de novo* assembled using default settings within CLC Genomics Workbench v9.5.2 (QIAGEN) with a minimum contig size threshold of 500 bp in length.

### Genome and plasmid annotations

We used RAST (http://rast.nmpdr.org/rast.cgi) for all annotations performed in this study (59).

### *in silico* serotyping

The serotype of each strain analyzed in this study was confirmed using the genes deposited in the Center for Genomic Epidemiology (http://www.genomicepidemiology.org) for *E. coli* as part of their web-based serotyping tool (SerotypeFinder 1.1 - https://cge.cbs.dtu.dk/services/SerotypeFinder) (60,60,61).

### *in silico* MLST phylogenetic analysis

The initial analysis and identification of the strains were performed using an *in silico E. coli* MLST approach, based on the information available at the *E. coli* MLST website Enterobase (http://enterobase.warwick.ac.uk/species/index/ecoli) and using Ridom SeqSphere+ software v2.4.0 (Ridom; Münster, Germany) (http://www.ridom.com/seqsphere). Seven housekeeping genes *(dnaE, gyrB, recA, dtdS, pntA, pyrC,* and *tnaA),* described previously for *E. coli* (37), were used for MLST analysis.

### *in silico* determination of virulence genes

Virulence genes were determined as previously described (9) using the genes deposited in the Center for Genomic Epidemiology (http://www.genomicepidemiology.org) for *E. coli* as part of their VirulenceFinder 1.5 web-based tool (https://cge.cbs.dtu.dk/services/VirulenceFinder) (61), We used Ridom to batch screen our set of genomes for known virulence genes. Supplementary Table 1 shows the 95 virulence genes analyzed by this method.

### *in silico* identification of antimicrobial resistance genes

We identified the antimicrobial resistance (AMR) genes present in our sequenced genomes as previously described by Gonzalez-Escalona (9), using genes deposited in the Center for Genomic Epidemiology (http://www.genomicepidemiology.org) as part of their web-based Resfinder 2.1 tool (https://cge.cbs.dtu.dk/services/ResFinder) (62). We used Ridom to batch screen our set of genomes for known AMR genes.

### Comparisons of genomic synteny

The genomic synteny between PacBio and MinION data was determined using Mauve (41).

### *stx* phage and T3SS identification and location

Prophages and prophage-like elements within the sequenced O26:H11 STEC strains were initially identified using the prophage-predicting PHASTER web server (40). Next, we used the genomic island prediction web server IslandViewer3 to detect potential pathogenicity and genomic islands (64). Each identified prophage, prophage-like element, and IE was then examined, using CLC Genomics Workbench 7.5, to locate nearby integrases and potential integration sites, which would confirm their status.

### Phylogenetic relationship of the strains by cgMLST analysis

Due to intrinsic problems with the sequencing technology used by MinION sequencing – generation of many indels located mainly in areas of homopolymers tracks-we wanted to test the effectiveness of MinION genomes produced by our pipeline to estimate the phylogeny of those 3 STEC strains in a context of 155 other genomes of highly similar O26:H11 available at NCBI. All these O26 genomes from GenBank were generated by MiSeq or HighSeq (Illuimina), considered the “gold standard” for accurate genome sequence determination and SNP analyses. We used Ridom SeqSphere+ software v2.4.0 to assess the phylogenetic relationship of these strains, performing a core genome multilocus sequence typing (cgMLST) analysis as previously described for O26:H11 (9). cgMLST uses the allele numbers of each loci to determine genetic distances and build the phylogenetic tree. We used O26:H11 strain 11368 (NC_013361.1) as the reference genome for generating the core genes for the phylogenetic tree and downloaded 195 genomes of E. coli O26:H11, available at NCBI, to build the phylogenetic tree. We used Nei’s DNA distance method (65) for calculating the matrix of genetic distance, taking only the number of same/different alleles in the core genes into consideration. A Neighbor-Joining (NJ) tree using the appropriate genetic distances was built after the cgMLST analysis.

### Nucleotide sequence accession numbers

The SRA sequences of all three *E. coli* strains used in our study are available in GenBank under the accession numbers: MiSeq data (SRR8333591, SRR8333592, and SRR8333590), MinION data (SRR8335317, SRR8335318, and SRR8335317), and PacBio data (xx, xx, and xxx).

